# *Pc*GCE is a potent elicitor of defense responses in aspen

**DOI:** 10.1101/2021.09.23.460629

**Authors:** Evgeniy N. Donev, Marta Derba-Maceluch, Xiao-Kun Liu, Henri Colyn Bwanika, Izabela Dobrowolska, Mohit Thapa, Joanna Leśniewska, Jan Šimura, Alex Yi-Lin Tsai, Dan Boström, Leszek A. Kleczkowski, Maria E. Eriksson, Karin Ljung, Emma R. Master, Ewa J. Mellerowicz

## Abstract

Using microbial enzymes in transgenesis is a powerful means to introduce new functionalities in plants. Glucuronoyl esterase (GCE) is a microbial enzyme hydrolyzing the ester bond between lignin and 4-*O*-methyl-α-D-glucuronic acid present as a side chain of glucuronoxylan. This bond mediates lignin-carbohydrate complex (LCC) formation, considered as crucial factor of lignocellulose recalcitrance to saccharification. Previous studies showed that hybrid aspen (*Populus tremula* L. x *tremuloides* Michx.) constitutively expressing *Phanerochaete carnosa* Burt GCE (*Pc*GCE) had better efficiency of cellulose-to-glucose conversion but were stunned and had lower cellulose content indicating that more studies are needed to design strategy for deployment of this enzyme *in planta*. Here we report that the transgenic plants exhibit premature leaf senescence, increased accumulation of calcium oxalate crystals, tyloses and necrotic lesions and have strongly activated immune defense reactions as revealed by their altered profiles of transcriptomes, metabolomes and hormones in the leaves. To elucidate if these effects are triggered by damage-associated molecular patterns (DAMPs) or by *Pc*GCE protein perceived as a pathogen-associated molecular pattern (PAMP), we ectopically expressed in aspen an enzymatically inactive *Pc*GCE^S217A^. The mutated *Pc*GCE induced similar growth retardation, leaf necrosis and premature senescence as the active one, providing evidence that *Pc*GCE protein is recognized as PAMP. Transcriptomics analysis of young expanding leaves of *35S:PcGCE* plants identified several candidates for receptors of *Pc*GCE, which were not expressed in developing wood tissues. Grafting experiments showed that *Pc*GCE transcripts are not cell-to-cell mobile and that *PcGCE* expressing leaves augment systemic responses. In agreement, expressing *Pc*GCE in developing wood by using the wood-specific promoter (WP), avoided all off-target effects. Moreover, *WP:PcGCE* lines grew better than control plants providing evidence that this strategy can be used in transgenic crops dedicated for biorefinery.

## Introduction

Saprophytic and pathogenic microbes living on lignocellulose developed specific ways of decomposing it. The repertoire of microbial enzymes has been greatly explored for enzymatic lignocellulose digestion *in vitro* (Thapa et al., 2020). Additionally, such enzymes, generally not found among native plant enzymes, have been successfully used in different crops for post-synthetic modification of cell wall (Brandon and Scheller, 2020). For example, expression of the ferulic acid esterase from *Aspergillus niger* van Thieghem (*An*FAE) in the forage crop tall fescue (*Festuca arundinacea* Schreb.), increased biomass digestibility by cellulases (de Buanafina et al., 2008; 2010). Xylan, which is a key factor of lignocellulosic biomass recalcitrance by reducing cellulose accessibility, has been a main target for plant cell wall modification for biorefinery (Donev et al., 2018). Expression of fungal acetyl xylan esterases from *A. niger* (*An*AXE1) or from *Hypocrea jecorina* Berkeley & Broome (formerly *Trichoderma reesei* Simmons) (*HjAXE*) in hybrid aspen (*Populus tremula* L. *x tremuloides* Michx.) increased yield of glucose by 20 to 30% in enzymatic saccharification without pretreatment with additional benefits of increased lignin solubility (Pawar et al., 2017b) and cell wall nanoporosity Wang et al., 2020). Even greater saccharification benefits were reported in poplar (*Populus alba* L.) expressing fungal xylanase *HvXYL1* (Kaida et al., 2009). Moreover, expressing microbial enzymes targeting the primary plant cell wall constituents, xyloglucan and pectins, also proved successful in reducing plant lignocellulose recalcitrance (Park et al., 2004; Kaida et al., 2009; Tomassetti et al., 2015).

However, microbial enzymes expressed *in planta* to modify plant cell wall sometimes triggered diverse immune responses, such as premature leaf senescence, necrotic lesions in the leaves, accumulation of reactive oxygen species (ROS) and induction of genes involved in biotic and abiotic stress responses (Bailey et al., 1990; Avni et al., 1994; de Buanafina 2012; Pogorelko et al., 2013; Tsai et al., 2017). It has been suggested that cell wall modifying enzymes could generate oligosaccharide fractions such as, oligogalacturonides (OGs) (Legendre et al., 1993; Norman et al., 1999; D’Ovidio et al., 2004), cellooligomers (Souza et al., 2017) or xyloglucan oligosaccharides (Claverie et al., 2018), which can be perceived by the plant cell as damage-associated molecular patterns (DAMPs).

Plants are also capable to recognize some exogenous microbial compounds, known as microbe- or pathogen-associated molecular patterns (MAMPs/PAMPs) (Boller and Felix 2009; Raaymakers and Van den Ackerveken 2016). For example, the 22-amino-acid N-terminal peptide of flagellin (flg22) of gram-negative bacteria displays a conserved elicitor activity across plant kingdom (Felix et al., 1999). Fragments of chitin, which is a fungal cell wall homopolymer shared among many classes of pathogens (Latgé, 2007), also act as PAMPs (Felix et al., 1998). Microbial enzymes can also be perceived as foreign compounds. For example, the ethylene-inducing xylanase (EIX) from *Trichoderma viride* Pers. has been discovered, which independently of its xylan degradation activity (Enkerli et al., 1999; Rotblat et al., 2002) induces pathogenesis-related (PR) proteins in tobacco (*Nicotiana tabacum* L.) and tomato plants (*Solanum lycopersicum* L.) (Bailey et al., 1990; Avni et al., 1994; Sussholz et al., 2020).

PAMPs and DAMPs are recognized by cell surface-localized receptor proteins, called pattern recognition receptors (PRRs) (Couto and Zipfel 2016; He et al., 2018). The stimulation of PRRs activates basal resistance response against broad range of pathogens, called pattern-triggered immunity (PTI) (Yu et al., 2017). Moreover, virulence factors could activate cytoplasmic intracellular resistance (R) proteins and initiate effector triggered immunity (ETI) (Jones and Dangl, 2006; Cui et al., 2015), which amplifies the basal PTI transcriptional program and triggers localized programmed cell death (PCD) (Balint-Kurti, 2019). Thus, the expression of microbial enzymes in plants potentially could trigger DAMP or PAMP signaling and activate defense responses and immunity. Indeed, transgenic plants expressing microbial enzymes frequently exhibit increased immunity against different pathogens (Pogorelko et al., 2013; Klose et al., 2015; Pawar et al., 2016; Reem et al., 2020).

Among different microbial enzymes, glucuronoyl esterase (GCE) has a potential for decreasing lignocellulose recalcitrance by hydrolyzing the ester bond between 4-*O*-methyl-α-D-glucuronic acid and lignin (Spániková and Biely, 2006; Biely et al., 2015; Bååth et al., 2016). This bond is thought to mediate the formation of lignin-carbohydrate complexes (LCCs) in woody species, considered as crucial factors of lignocellulose recalcitrance (Giummarella et al., 2019). When GCE from necrotrophic wood decaying white-rot basidiomycete, *Phanerochaete carnosa* Burt (*Pc*GCE), was ectopically expressed in *Arabidopsis thaliana* (L.) Heynh. and hybrid aspen, leaf senescence was accelerated and growth was reduced (Tsai et al., 2012; Gandla et al., 2015; Tsai et al., 2017). The wood of transgenic hybrid aspen had increased lignin and decreased cellulose and extractives contents (Gandla et al., 2015). But despite highly elevated lignin content, the conversion of wood cellulose to glucose (Glc) after acid pretreatment was increased as predicted. To elucidate the cause of untargeted effects of *Pc*GCE in transgenic plants, we characterized physiological responses to the transgene in more detail, determining changes in their transcriptome, metabolome, in levels of their hormones and reactive oxygen species (ROS). We also investigated the progress of these changes during leaf development and their transmission through grafts. The results indicated that *Pc*GCE is a potent elicitor of stress responses in aspen recognized by PAMP signaling, most probably in the leaves. Moreover, we demonstrated that expressing the same enzyme using the wood-specific promoter, avoids all undesirable responses providing a strategy for deployment of this enzyme in transgenic crops dedicated for biorefinery. The analysis also revealed novel genes and metabolic pathways in aspen involved in perception of pathogens and in downstream defense reactions.

## Results

### Ectopically expressed Pc*GCE* induced developmental defects and defense responses in aspen

Hybrid aspen lines ectopically expressing *Pc*GCE displayed reduced stem height and diameter, and root dry weight compared to wild-type (WT) (Fig. 1, ABC). As observed previously, all transgenic lines developed leaf necrosis and premature senescence (Gandla et al., 2015). The necrotic spots first appeared on fully expanded leaves, then they rapidly enlarged occupying most of the leaf area, (Fig. 1D) and the leaves were shed prematurely. Using a dye uptake analysis, we found that the hydraulic continuity was compromised in the leaves of transgenic plants before the necrotic spots appeared (Fig.1, EF). This was visible as unstained areas on leaves of branches immersed in a staining solution. Anatomical analyses of the unstained areas revealed that xylem vessels were blocked by gels and tyloses (Fig. 1, GH), indicative of activation of defense responses to pathogen attack and/or herbivory and signaling by ethylene (ET) and jasmonic acid (JA) (Leśniewska et al., 2017).

**Figure 1.**
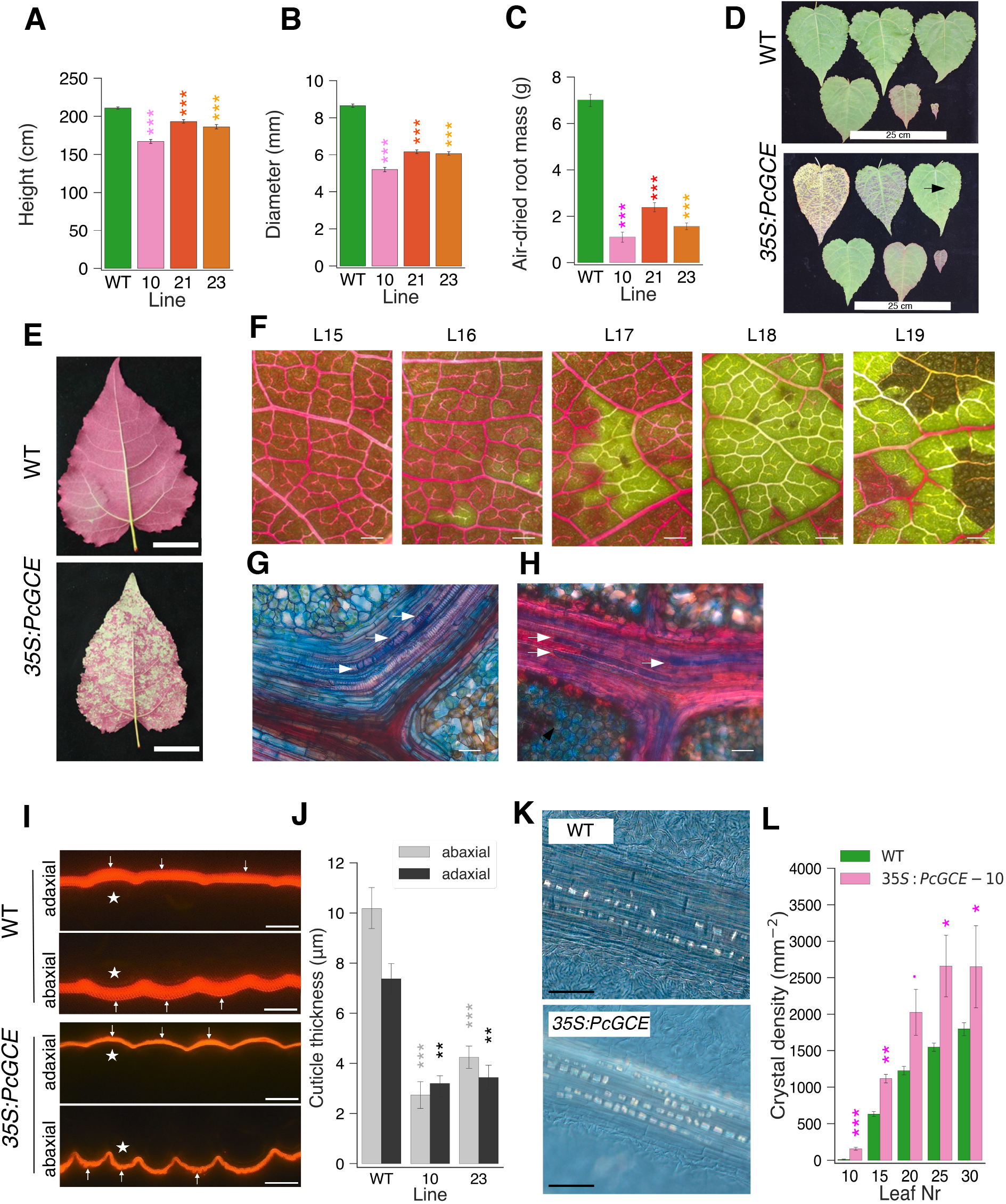
Transgenic aspen constitutively expressing *PcCGE* display different disease symptoms. Height (A), diameter (B), and root weight (C) of eleven-week-old plants of three independent transgenic lines: 10, 21 and 23 and WT. *N*=24 for transgenic and 40 for WT plants in (A) and (B), and 6 plants in (C). (D) Appearance of developing leaves in transgenic and WT plants. Note the rapid progression of necroses in leaves of the transgenic plant. Shown are leaves 5, 10, 15, 20, 25 and 30 of line 10 and WT plants. First necrotic spots are pointed by the arrow. (E, F) Hydraulic continuity is compromised in transgenic leaves as shown by the dye uptake inhibition. Green areas indicate hydraulic blockage. (E) Leaf 22 in transgenic (line 10) and WT plants. Scale bar = 5 cm. (F) Progression of dye uptake blockage and necroses in successive leaves of a line 10 plant. Scale bar = 500 μm. (G, H) Blockage of xylem vessels by tyloses (G, arrows) and gels (H, arrows) in the leaves of transgenic plants (line 10). Longitudinal sections through vascular bundles stained with safranin and alcian blue. Scale bars = 20 μm. (I) Cuticle (arrows) in leaf epidermis of line 10 and WT plants stained with Nile red. * is placed over epidermal cells, scale bars: 10 μm. (J) Cuticle thickness in transgenic lines and WT. (K) Calcium oxalate crystals in the leaf veins of line 10 and WT plants. Note the higher density and larger size of crystals in transgenic plants. Scale bar = 100 μm. (L) Crystal density over veins in leaves 10-30 of line 10 and WT plants. Asterisks show significant differences compared to WT (Student’s T-test). Statistical significance: · - P< 0.1; * - *P* ≤ 0.05; ** - *P* ≤ 0.01;*** - *P* ≤ 0.001.

To see if other defense responses were activated in transgenic plants, we analyzed cuticle and lipids which are known to respond to biotic stresses (Ziv et al., 2018). The abaxial and adaxial epidermis of transgenic plants exhibited strongly reduced cuticle thickness (Fig. 1, IJ). This was accompanied by a reduction of several species of free fatty acids (FAs) as evidenced by analysis of leaves 10, 13, and 15 in line 10 and in WT. In young expanding leaves (leaf 10), the FAs with chains of 18 carbons or longer, were affected including C18:1 cis 9, C18:2 cis 9 and 12, C20:0, C21:0 and C22:0 (Fig. S1, AB). In older fully expanded leaves (leaf 13 and 15), the changes were more pronounced and C18:0 FA had also reduced content in transgenic plants. These results indicate a disruption of very long chain fatty acids (VLCFA) biosynthesis in transgenic plants. Since VLCFA are essential for wax biosynthesis, we examined wax profiles in chloroform extracts of expanded leaves of line 10, line 23 and WT plants using GC-MS. Unexpectedly, the transgenic plants showed increased contents of chloroform-extracted fatty wax components, including mid-chain and long-chain FA, long-chain fatty esters, especially C37:3 ester, fatty alcohols, and alkanes, and glycerol, whereas the content of eluted cinnamic acid was greatly reduced (Fig. S1C). Overall, the data show that the transgenic plants exhibit disrupted long- and VLCFA biosynthesis, thinner cuticle and that they have altered composition of chloroform-extracted wax compounds.

Herbivory stress is known to induce the accumulation of calcium oxalate (CaOx) crystals (Molano-Flores, 2001). Therefore, we investigated if the abundance of crystals was increased in leaves of transgenic plants. In hybrid aspen, prismatic crystals are localized mostly along the veins and their density increases with leaf age (Fig. 1, KL, Fig. S2). The crystals displayed higher density and larger size in transgenic plants compared to WT plants. The chemical identity of the crystals was established by X-ray diffraction as calcium oxalate (CaOx) or whewellite, CaC_2_O_4_ x H_2_O.

In all, the above data suggested that aspen plants constitutively expressing *Pc*GCE activate defense responses to herbivory and pathogen attack. Since ROS induction is one of first responses following pathogen attack, we examined ROS levels in fully expanded leaves 21 and 23 of the three transgenic lines, 10, 21 and 23 by diaminobenzidine (DAB) staining (Fig. 2, AB). Leaf 21 was always symptom-free, whereas leaf 23 typically exhibited first necrotic spots. Both leaves of all three transgenic lines showed a strong increase in DAB staining compared to WT, and an increase in staining in the older compared to younger leaf, indicative of increased ROS levels.

**Figure 2.**
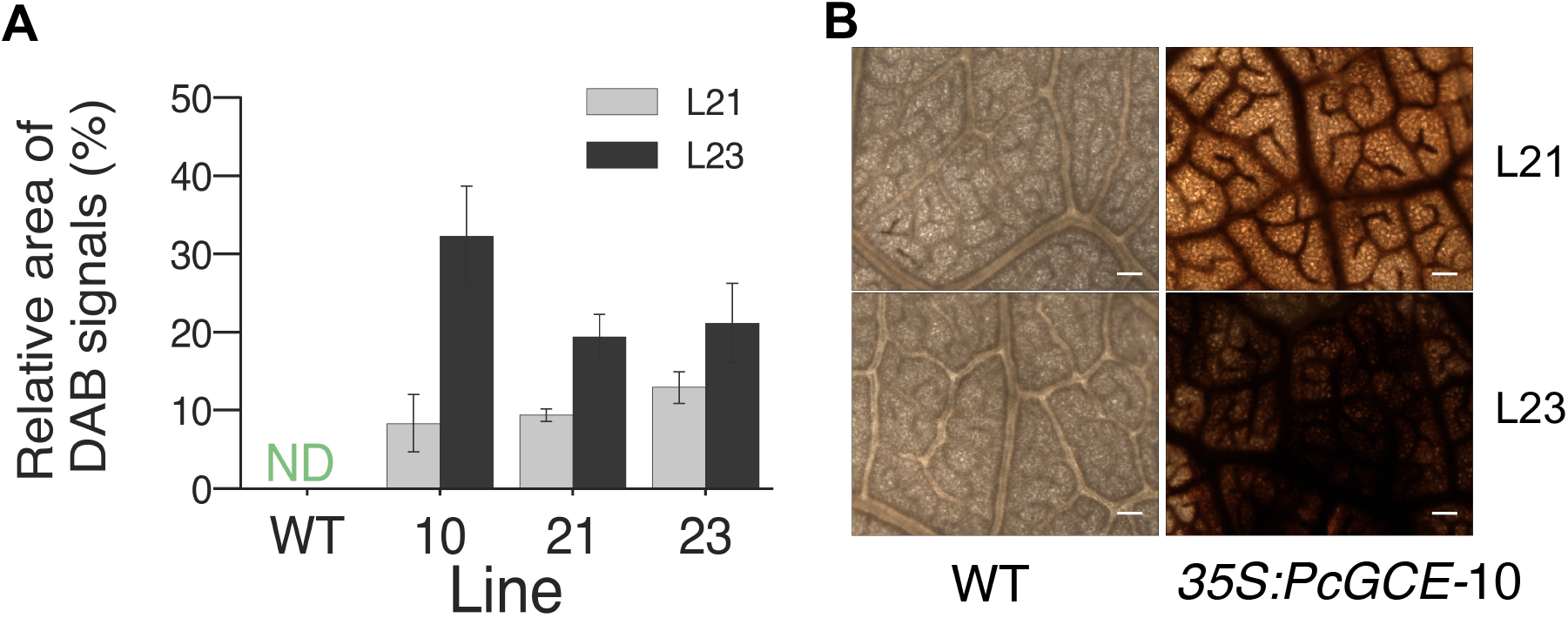
*Pc*GCE induces ROS accumulation in the leaves of *PcGCE* expressing plants. Symptomless leaves (L21) and in leaves with first necroses (L23) were analyzed by diaminobenzidine (DAB) staining in three independent lines carrying *35S:PcGCE*: 10, 21. and 23. (A) Area of DAB signals (pixels with intensity between 0 and 31) quantified by image analysis. Means ± SE, n = 12. DAB signals were not detected (ND) in WT. (B) Light microscopy images of representative leaf 21 and 23 (L21 and L23) in transgenic line *35S:PcGCE-10* and WT. Scale bar = 100 μm.

### Molecular changes caused by Pc*GCE* in aspen leaves

To characterize the molecular pathways used by *Pc*GCE to stimulate variety of stress symptoms observed (Figs. 1, 2), we analyzed transcriptomes, metabolomes and hormonomes of leaves 21 and 23 in transgenic lines 10 and 23, and in WT. We found that the lines expressing *Pc*GCE had massively disrupted transcriptome profiles with as many as 16,087 genes (39% of all protein-coding genes) differentially expressed (DE) in at least one line and at least in one leaf developmental stage (Table S2), and 4,143 and 6,126 genes (10 and 15% of protein coding genes) differentially expressed in leaf 21 and leaf 23, respectively, in both transgenic lines (Fig. 3A, Table S3). The gene ontology analysis of DE genes in both lines (Fig. 3B) showed enrichment in oxidation-reduction processes, regulation of transcription, defense signaling in biotic and abiotic stresses, signaling by JA, SA and ET, metabolic processes, response to red light, malate metabolism, and photosynthesis. The biotic stress related genes affected in both lines (233 genes) were mostly upregulated, in agreement with the physiological changes observed in the transgenic plants, whereas abiotic response genes (143 genes) were mostly downregulated (Table S3). Among redox-related genes (107) there were 21 peroxidases, which were all upregulated. Among DE phenyl propanoid (49 genes) and flavonoid biosynthetic (36 genes), pathway genes the majority were highly upregulated. Almost all JA-related DE genes (51) were upregulated, whereas SA-related DE genes (18) were mostly downregulated. ET-related DE genes (105) were strongly either up- or downregulated. Among 118 photosynthesis-related DE genes, the majority were downregulated. Among a large group of receptor-encoding DE genes (453), majority were upregulated, particularly in the younger leaf. Among 365 identified DE genes encoding channels and transporters, majority of genes encoding ABC, and sulphate transporters were upregulated, whereas those encoding aminoacidic transporters were downregulated. The latter could be related to general downregulation of genes involved in protein biosynthesis in transgenic lines. We identified 517 putative transcription factors (TFs) among the DE genes, majority of which were downregulated, which corresponded to downregulation of majority of genes involved in developmental programs, DNA and chromatin organization, and transcription. Opposite to this general trend, several MYB TFs, including MYB15, MYB62, MYB63, MYB66, MYB68, MYB73, MYB84, and MYB116, and several WRKY TFs, such as WRKY6, WRKY51 and WRKY75 were highly upregulated.

**Figure 3.**
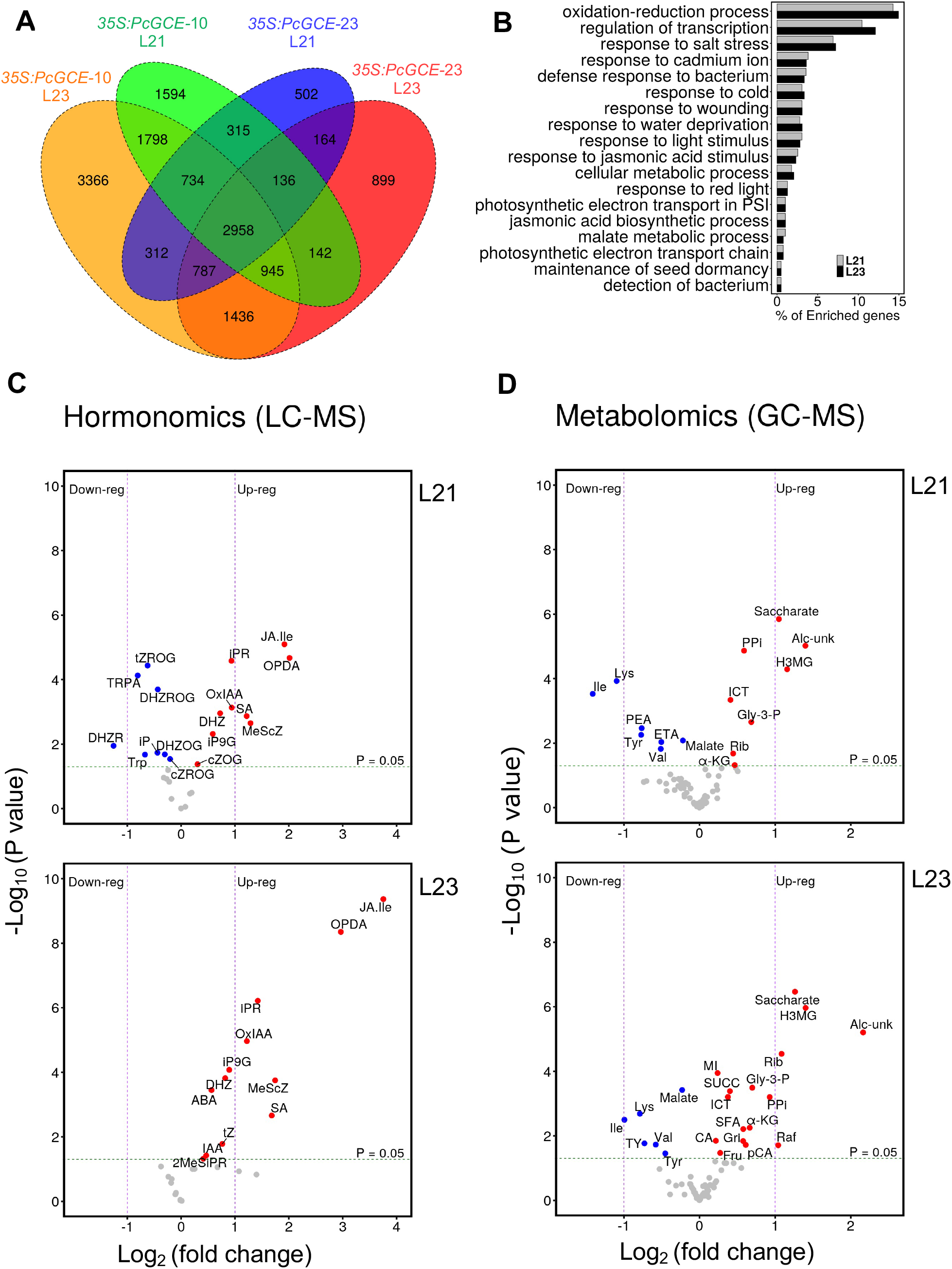
Overview of transcriptomic and metabolomic changes in aspen ectopically expressing *PcGCE*, compared to wild type. Symptomless leaves (L21) and in leaves with necrosis (L23) were analyzed in two independent transgenic lines: *35S:GCE-10* and *35S:GCE*-23. (A) Venn diagram of differentially expressed genes (DEGs). (B) Bar graph of Gene Ontology (GO) enrichment analysis of DEGs in both transgenic lines. (C, D) Volcano plots showing altered levels of metabolites detected by hormonomics (C) and metabolomics (D) analyses in leaves 21 and 23. The significantly altered metabolites compared to WT are labeled (Student’s T-test). **Hormone abbreviations (C):** ABA – abscisic acid; cZOG – cis-zeatin-O-glucoside; cZROG – cis-zeatin riboside-O-glucoside; DHZ – dihydrozeatin; DHZR – dihydrozeatin riboside; DHZOG – dihydrozeatin-O-glucoside; DHZROG – dihydrozeatin riboside-O-glucoside; iP – N6-isopentenyl-adenine; iPR – N6-iso-pentenyl-adenenosine; JA – jasmonic acid; JA-Ile; MeScZ – 2-methylthio-cis-zeatin; MeSiPR – 2-methylthio-isopentenyladenosine; IAA – Indole-3-acetic acid, iP9G – N6-isopentenyladenine-9-glucoside; OPDA – cis-12-oxo-phytodienoic acid; OxIAA - 2-oxoindole-3-acetic acid; SA – Salicylic acid; Trp – tryptophan; TRPA – Tryptamine; tZ – trans-zeatin, tZROG – trans-zeatin-riboside-O-glucoside. **Metabolite abbreviations (D):** Alc-unk – unknown alcohol; CA – caffeate; ETA – EtOH-amine; Fru – fructose; Gly-3-P – glycerol-3-P; Gri – glyceric acid; H3MG – hydroxy-3-methylglutaric acid; ICT – isocitrate; Ile – iso-leucine; α-KG – keto-glutarate; Lys – lysine; MI – *myo*-inositol; pCA – *p*-coumarate; PEA – phenetyl-amine; PPi – pyro-phosphate; Rib – ribose; SUCC - succinate; SFA – sulfamic acid; TY – tyramine; Val – valine.

The hormone profiling (Šimura et al., 2018) further provides evidence of hormonal signaling of stress responses downstream of *Pc*GCE. The analysis targeted compounds related to cytokinins, auxins, gibberellins, jasmonates, SA and brassinosteroids, but not ET. Leaf 21 hormone profile (Fig. 3C) showed induction of JA and its precursor 12-oxophytodienoic acid (OPDA), SA, inactivated auxin (OxIAA), cytokinin precursor isopentenyl-adenenosine (IPR), active cytokinin dihydrozeatin (DHZ) and some of their degradation products [cis-zeatin-O-glucoside (cZOG), isopentenyladenine-9-glucoside (IP9G)], whereas an active cytokinin isopentenyl-adenine (IP) and cytokinin degradation products [dihydrozeatin riboside-O-glucoside (DHZROG), dihydrozeatin riboside (DHZR), dihydrozeatin-O-glucoside (DHZOG), cis-zeatin riboside-O-glucoside (cZROG), trans-zeatin-riboside-O-glucoside (tZROG)] were reduced. Furthermore, auxin precursors, tryptamine (TRPA) and tryptophan (Trp) were downregulated. In leaf 23, we observed increased levels of hormones affected in the younger leaf and in addition, an induction of SA, ABA, IAA and active cytokinins [DHZ and trans-zeatin (tZ)] cytokinin precursor IPR as well as cytokinin degradation product IPA9G and cytokinins with unknown function (methylthio-cis-zeatin (MScZ) and methylthio-isopentenyladenosine (MeSiPR)], which correlated with the formation of leaf necrotic spots. Thus, the analysis revealed altered hormonal levels of stress-related phytohormones and cytokinins, which are considered to be a hub for defense responses mediated by other hormones (Choi et al. 2011).

Metabolomic analysis (Fig. 3D), revealed upregulation of hydroxy-3-methyl glutaric acid (mevalonate), myo-inositol, glycerol-3-phosphate (Gly-3-P), saccharate, ribose, and metabolites related to the TCA cycle, such as iso-citrate, and keto-glutarate. In contrast, the contents of aminoacids (Ile, Lys, Tyr, Val) and malate were reduced. Contents of fructose, raffinose, succinate, *p*-coumaric acid and caffeic acid were upregulated in leaf 23 of transgenic plants, underlying the progressive response to transgene in older leaves. Mevalonate (Nelson et al., 1994) and Gly-3-P (Chanda et al., 2008) have been associated with basal resistance and induction of systemic immunity. Furthermore, enhanced levels of sugars have been observed during stress response (Gómez-Ariza et al., 2007; Conrath, 2011) and may indicate formation of sink at the pathogen perception site (Sutton et al., 2007; Essmann et al., 2008). Lower amino acid content corresponds to general decrease in protein biosynthesis and amino acid metabolism reflected in transcriptome (Table S3).

Overall, the omics analyses revealed heavily disrupted metabolism in transgenic plants, with activation of biotic stress and downregulation of growth and development.

### Sequential analysis of developing leaves revealed three distinct stages of disease development

Since transcriptomics, hormonomics and metabolomics analyses revealed more differences between transgenic and WT plants in leaf 23 than in leaf 21, we investigated if the expression of transgene is stable in sequential leaf samples from the apical bud to mature necrotic leaves by reverse transcription quantitative polymerase chain reaction (RT-qPCR) analysis. It showed that *Pc*GCE mRNA was accumulating exponentially in developmental leaf series (Fig. 4A).

**Figure 4.**
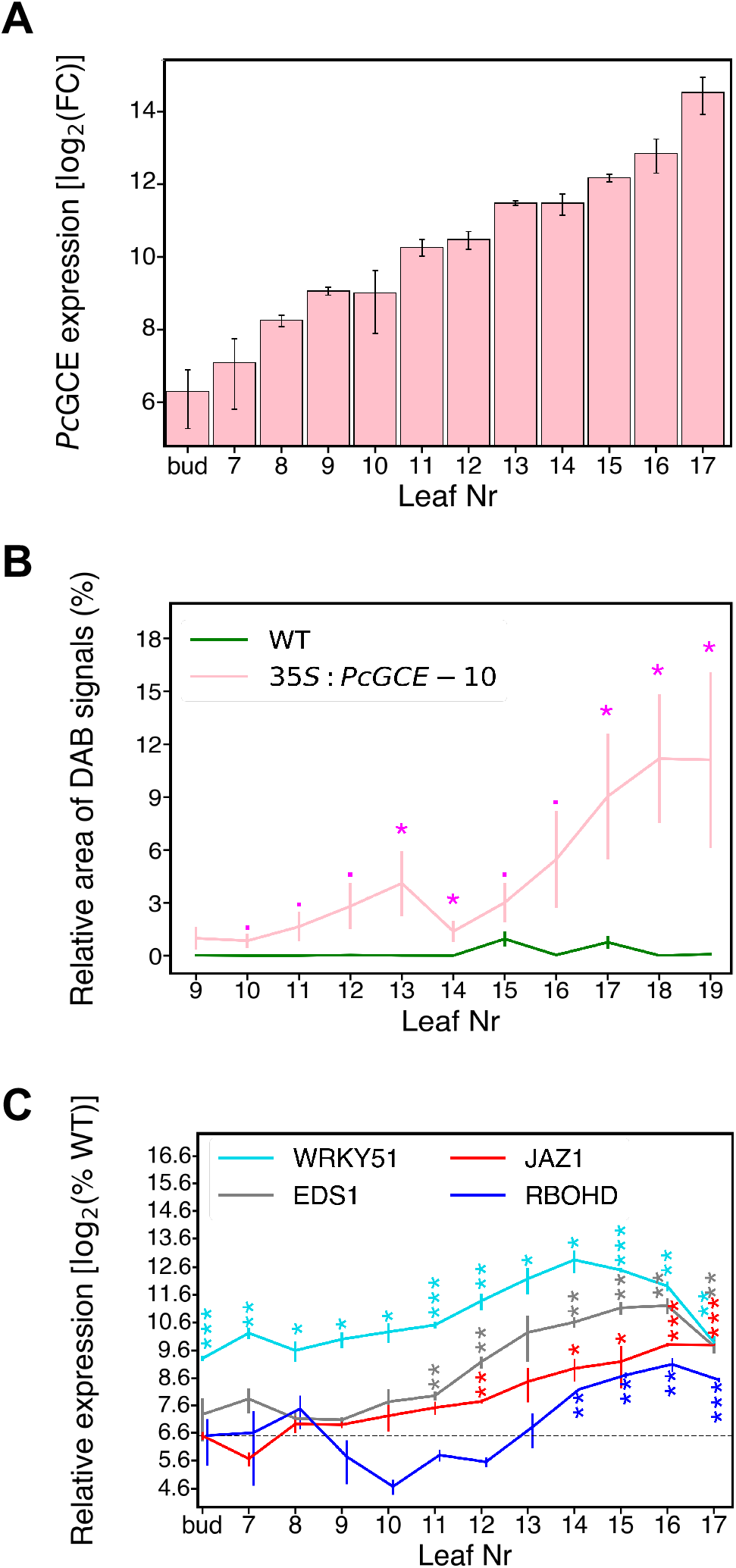
Expression of *PcGCE* and development of defense reactions in sequential leaves of 8-week-old plants of transgenic line 10. (A) *PcGCE* transcript levels in developing leaves relative to levels in the bud based on RT-qPCR. (B) ROS levels analyzed by DAB staining. DAB signals were pixels with gray scale level between 0 and 31. (C) Transcript levels of defense response marker genes: *JAZ1* (*Potri.003G068900*), *RBOHD* (*Potri.001G070900*), and stress marker genes involved in response to low oleic acid levels *WRKY51* (*Potri.005G085200*) and *EDS1* (*Potri. 015G069600*). Data are means ± SE, n=3 biological replicates in A and C or 12 replicates in B. Asterisks show significant differences compared to WT (Student’s test; · - P≤ 0.1; * - P ≤ 0.05; ** - P ≤ 0.01; *** - P ≤ 0.001).

We further investigated the pattern of defense responses in successive leaf samples. ROS content in transgenic plants showed an early small peak around leaf 13, and then a rapid increase from leaf 14 to 18 (Fig. 4B). No DAB signals were seen in WT. We further studied expression of marker genes selected among the DE genes in leaves 21 and 23 (Fig. 4C). The selected genes represented different pathways based on the affected processes as revealed by transcriptomics. Homolog of *JAZ1* (*Potri.003G068900*) representing JA signaling pathway responsive to JA levels (Chung et al., 2008) was highly upregulated starting from leaf 8 and further increasing up to 16-fold in leaf 17. The high expression was maintained in leaves 21 and 23 (Table S3). Homolog of *RBOHD* (*Potri.001G070900*) representing ROS generating pathway of systemic acquired resistance (SAR) triggering hypersensitive response (HR) response (Nühse et al., 2007; Lew et al., 2020) was upregulated only in older, necrosis-affected leaves (Fig. 4C), corresponding to the main ROS peak (Fig.4B). Also, homologs of *EDS1* (*Potri.015G069600*) and *WRKY51* (*Potri.005G085200*), involved in HR and antagonizing JA signaling (Gao et al., 2011) were induced at high levels in older leaves (Fig. 4C). Thus, the analysis revealed two main stages in the progress of *Pc*GCE stress induction in developing leaves, namely an early stage dominated by JA signaling, and a late stage characterized by SAR driven HR.

### Grafting experiments revealed that JAZ1 and RBOHD were induced at a distance from the stimulus but the transgene mRNA was not mobile

Since the above transcriptomics (Table S3), hormonomics (Fig. 3C) and sequential leaf analyzes (Fig. 4) demonstrated upregulation of signaling by JA, SA, as well as ROS accumulation and SAR development, we wondered which of defense reactions were systemically induced by *Pc*GCE. To distinguish between local and systemic responses to *Pc*GCE, the scions of five-week-old transgenic and WT plants were grafted onto the transgenic or WT rootstocks in all possible combinations (Fig. 5A). First, we analyzed the presence of *Pc*GCE transcripts in the grafted plants and found them only in the leaves of transgenic scions and rootstocks (Fig. 5B). This indicates that the transcripts are not transmitted to WT shoots or rootstocks of the grafted plants, and thus any disease symptoms in non-transgenic parts must be induced by *Pc*GCE at a distance.

**Figure 5.**
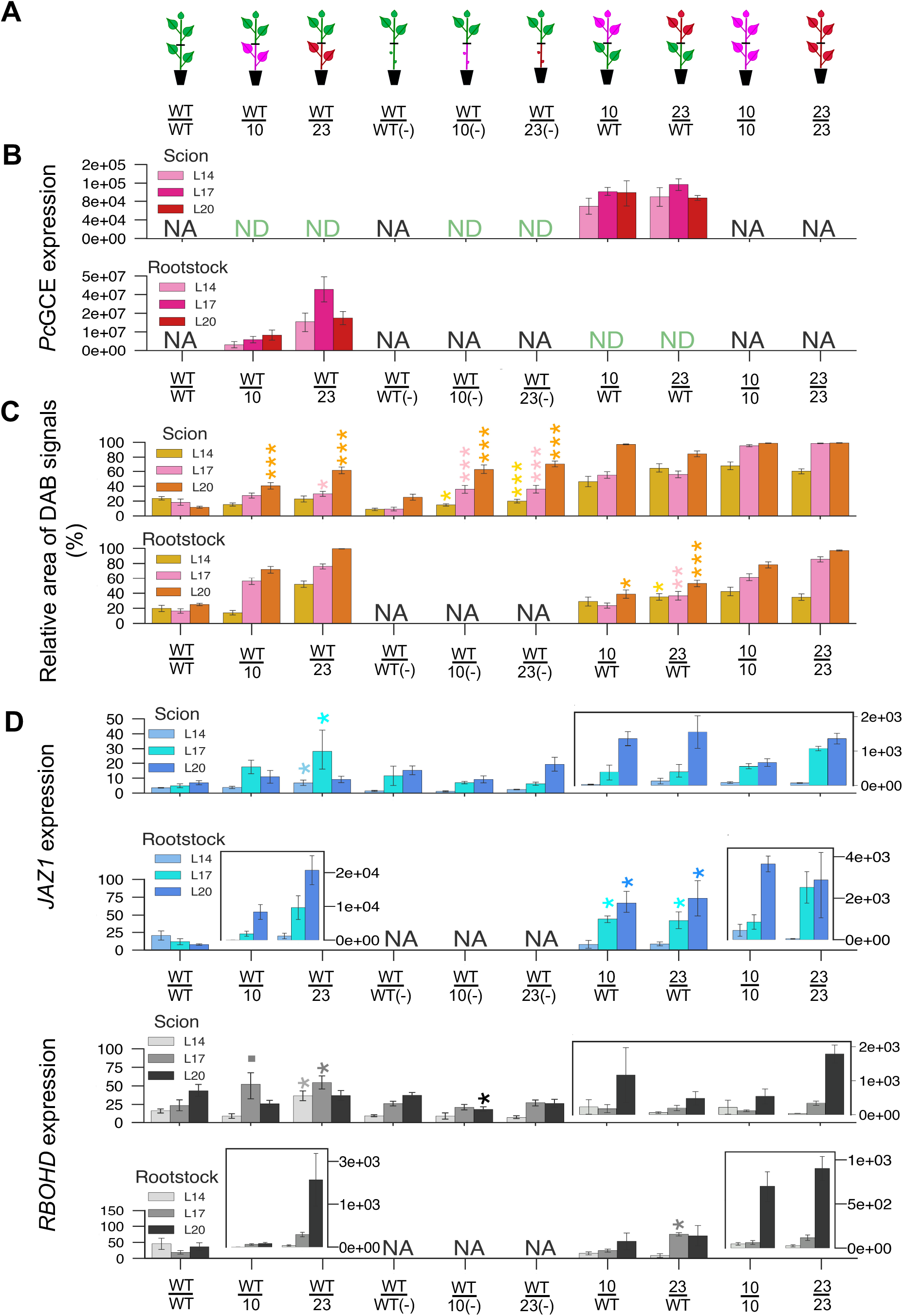
Grafting experiments indicate that defense responses are induced by *Pc*GCE at a distance. (A) Schematics of different types of grafts between WT and transgenic plants of line 10 and line 23. (-) indicates rootstocks where leaves were removed from the main stem. (B) Levels of *PcGCE* transcripts in leaves of scions and rootstocks. (C) ROS levels in leaves of scions and rootstocks determined by DAB staining and image analysis. (D) Marker gene induction in organs that do not express *PcGCE*. Leaves 14, 17 and 20 (L14, L17, L20) were analyzed 48 days after grafting. ND – not detected; NA – not analyzed. Expression values are fold change relative to the least expressing sample. Asterisks show significant differences compared to WT/WT (Student’s test, - P≤ 0.1; * - P ≤ 0.05; ** - P ≤ 0.01; *** - P ≤ 0.001). The statistical analysis is shown only for WT scions or rootstocks combined with transgenic rootstocks and scions, respectively.

Increase in ROS was observed in WT scions grafted on transgenic rootstocks, and in WT rootstocks carrying transgenic scions compared to WT/WT grafts (Fig. 5C), indicating that this SAR symptom is graft-transmittable and induced at a distance by *Pc*GCE. Furthermore, removal of leaves from transgenic rootstocks did not affect ROS propagation to WT scions.

Expression of *JAZ1* and *RBOHD* was induced in leaves of WT scions grafted on transgenic rootstocks and in leaves of WT rootstocks carrying transgenic scions compared to WT/WT grafts (Fig. 5D). The induction of these genes in WT scions depended on the presence of transgenic leaves in transgenic rootstocks, suggesting that the transgenic leaves are important for the propagation of systemic signal responsible for marker gene induction.

### Expressing of Pc*GCE* from wood-specific promoter allows normal growth

The grafting experiments showed that although stress symptoms and marker genes were induced at a distance when *Pc*GCE was expressed in distant plant organs, the transcripts of the transgene were not mobile in the plant. This suggests that expressing *Pc*GCE in a tissue that is unable to perceive this enzyme, would avoid development of stress symptoms. To test this hypothesis, we expressed the transgene in cells developing secondary cell walls using the wood-specific promoter (WP) (Ratke et al., 2015). Three lines most highly expressing *Pc*GCE (line 8, 9 and 14) were selected from 20 independent transformation events and grown in the greenhouse for 11 weeks. The lines did not show any growth defects and they grew slightly better than WT (Fig. 6A). In contrast, *35S:PcGCE* lines that were grown along exhibited pronounced premature leaf senescence and necrosis. Two lines of *WP:PcGCE* and *35S:PcGCE* constructs were subsequently selected for more thorough analyses. In the leaves, transcripts levels of *WP* lines were much reduced compared to *35S* lines whereas in developing wood the two types of transgenic plants exhibited comparable transgene expression (Fig. 6B). Growth analyses showed a significant increase in plant height and diameter in both transgenic *WP:PcGCE* lines, and an increase in root mass and leaf size in line 14 compared to WT (Fig. 6 CDEF). To check if using *WP* prevented ROS accumulation in the leaves of transgenic lines, we analyzed leaves at successive stages of development by DAB staining. Interestingly, ROS levels were slightly higher compared to WT, especially in line 8 that was more highly expressing the transgene (Fig. 6GH), but in contrast to *35S:PcGCE* lines, which exhibited progressive increase in ROS during leaf development (Fig. 3B), the levels of ROS in *WP:PcGCE* lines remained stable (Fig. 6G).

**Figure 6.**
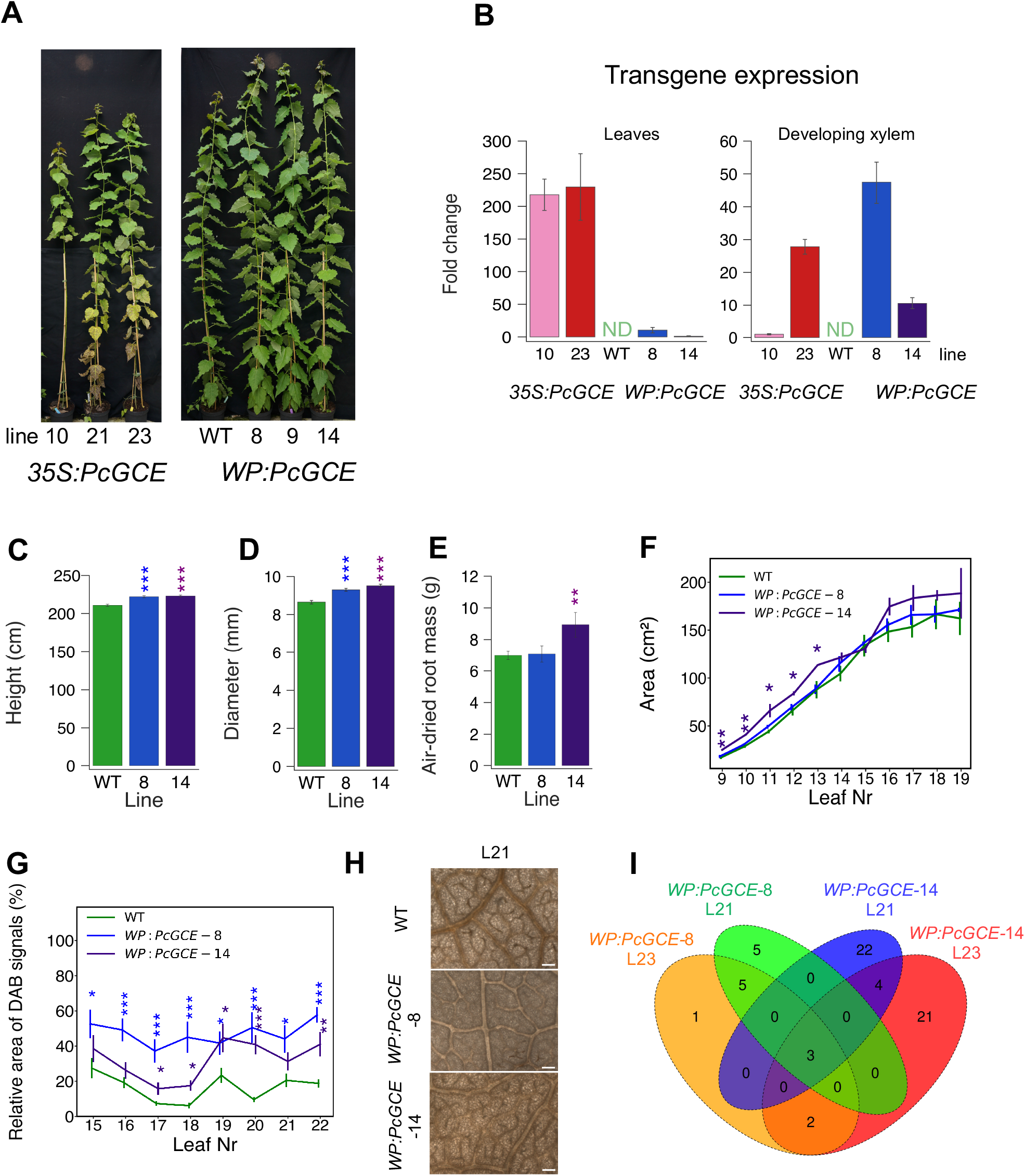
Transgenic aspen expressing *PcGCE* from the *WP* promoter does not show disease symptoms during the 11-week-cultivation in the greenhouse. (A) Appearance of *WP:PcGCE*-8, *WP:PcGCE*-9 and *WP:PcGCE*-14 lines compared to WT trees and lines expressing *PcGCE* constitutively. (B) Transgene expression in mature leaves (L21) and developing xylem of three biological replicates of *35S:PcGCE* and *WP:PcGCE* lines analyzed by RT-qPCR. No expression was detected in WT plants (ND). Expression was normalized to the lowest-expressing line. Height (C), diameter (D), dry root mass (E) and leaf area (F) of eleven-week-old plants. (G) ROS levels in leaves analyzed by DAB staining and image analysis. (H) Representative images of DAB-stained leaf 22 (L22) in transgenic and WT plants. Scale bar =100 μm. Data in B-G are means ± SE. Asterisks show significant differences compared to WT (Dunnett’s test, · - P< 0.1; * - *P* ≤ 0.05; ** - *P* ≤ 0.01;*** - *P* ≤ 0.001). (I) Venn diagram of differentially expressed genes (DEGs) in mature leaves 21 (L21) and 23 (L23) of transgenic lines: *WP:PcGCE*-8 and *WP:PcGCE*-14 compared to WT.

Transcriptomic analysis of the leaves revealed 62 DE genes as compared to WT in one or both transgenic lines in leaf 21 or 23 (Fig. 6I, Table S4). Only three of these genes were DE in both investigated lines in leaf 21, and these three genes were also affected in leaf 23. These genes included an extracellular cellulase *PtCel9B3* (Takahashi et al., 2009) which was upregulated, and two downregulated genes: a homolog of phytosulfokine receptor 1 (*PSKR1*) and a homolog of *MILDEW RESISTANCE LOCUS O 5* (*MLO5*), both involved in Ca-signaling during developmental and biotic stress responses (Mosher et al., 2013; Hartmann et al., 2014; Meng et al., 2020). Leaf 23 exhibited one more DE gene encoding histone 2A, which was upregulated (Table S4).

Hormonomics analysis revealed downregulation of active cytokinins, their degradation products, ABA and SA, and auxin precursor tryptamine in transgenic lines (Table S5). Cytokinin species reacted the same way in *WP:PcGCE* as in *35S:PcGCE* lines but with much reduced intensity. Other hormonal responses, particularly those concerning jasmonates and SA, which were strongly upregulated in *35S:PcGCE* lines (Fig. 3C), did not show similar response in *WP:PcGCE* plants (Table S5). Metabolomic analysis revealed slight upregulation of some monosaccharides (Table S6), which were seen strongly upregulated in *35S:PcGCE* lines (Fig. 3D). However, metabolites related to the TCA cycle, inorganic phosphate, caffeic acid, and amino acids were not altered in *WP:PcGCE* lines, suggesting that the activated metabolic pathways in lines with *35S* and *WP* promoters are not overlapping. These data clearly show that the growth inhibition and stress-related symptoms that were observed in *35S:PcGCE* expressing plants were avoided when the same protein was expressed from *WP* promoter.

### Transcriptomics of early stages of leaf development identified putative receptors and signaling components involved in the perception of Pc*GCE*

Since grafting experiments indicated that leaves could be the site of *Pc*GCE perception and source of signal that induces stress responses systemically, we analyzed transcriptomes of developing leaves in *35S:PcGCE* line 10 and WT plants at a younger stage to reduce influence of secondary effects of *PcGCE* expression and reveal candidate genes encoding proteins involved in *Pc*GCE perception. We targeted the earliest stages of leaf development to minimize secondary effects: the first unfolded leaf (leaf 8) and a young expanding leaf (leaf 11). Expression of the transgene was approximately two times higher in leaf 11 than in leaf 8 (Fig. 7A), confirming the RT-qPCR analysis (Fig. 4A), whereas the corresponding increase in the number of DE genes was approximately 4-fold, from 1,571 DE genes in leaf 8 to 6,328 DE genes in leaf 11, among which 941 genes were DE in both leaves (Fig. 7B; Table S7). This indicates that *Pc*GCE initially affects relatively small number of genes that in turn induce a cascade of transcriptomic responses. Gene ontology (GO) enrichment analysis of *Arabidopsis* homologous genes (Table S8) shows 65 significantly affected processes in leaf 8, including developmental processes as well as biotic and abiotic stress responses. Fewer and not much overlapping processes were affected in leaf 11. Functional annotation of *P. trichocarpa* DE transcripts revealed that the house-keeping activities and development were massively affected in leaf 11, whereas the response in leaf 8 was more attenuated. For example, genes involved in protein biosynthetic machinery, cell cycle, general transcriptomic machinery, DNA and chromatin organization and cytoskeleton were largely affected (downregulated) in leaf 11 but they were little altered in leaf 8 (Fig. 7C; Table S7). Moreover, these house-keeping and development-related categories had larger proportions of upregulated genes in leaf 8 as compared to leaf 11. In comparison, proportionally more genes were already affected in leaf 8 in categories “Receptors” and “Ca signaling” and these categories tended to be more abundantly represented in leaf 11. The genes of these categories were mostly upregulated. Relatively larger proportions of genes uniquely affected in leaf 8 were found for the “Photosynthesis” and “Starch and sugars metabolism”. These genes were also mostly upregulated at both leaf developmental stages but more so at a younger stage and there was relatively little overlap between the stages. “Cell wall polysaccharides” related genes were by large downregulated in leaf 8 and this trend tended to be reversed in leaf 11. Many lipid- and TCA metabolism-related genes were affected already in leaf 8. Thus, the earliest responses recorded in transcriptomes of developing leaves of transgenic plants were different from those seen at later stages, and indicated activation of biosynthesis of photosynthetic machinery, and induction of different receptors and proteins involved in Ca signaling. These early responses were followed by general downregulation of housekeeping activities and development in older leaves, and further development of stress responses.

**Figure 7.**
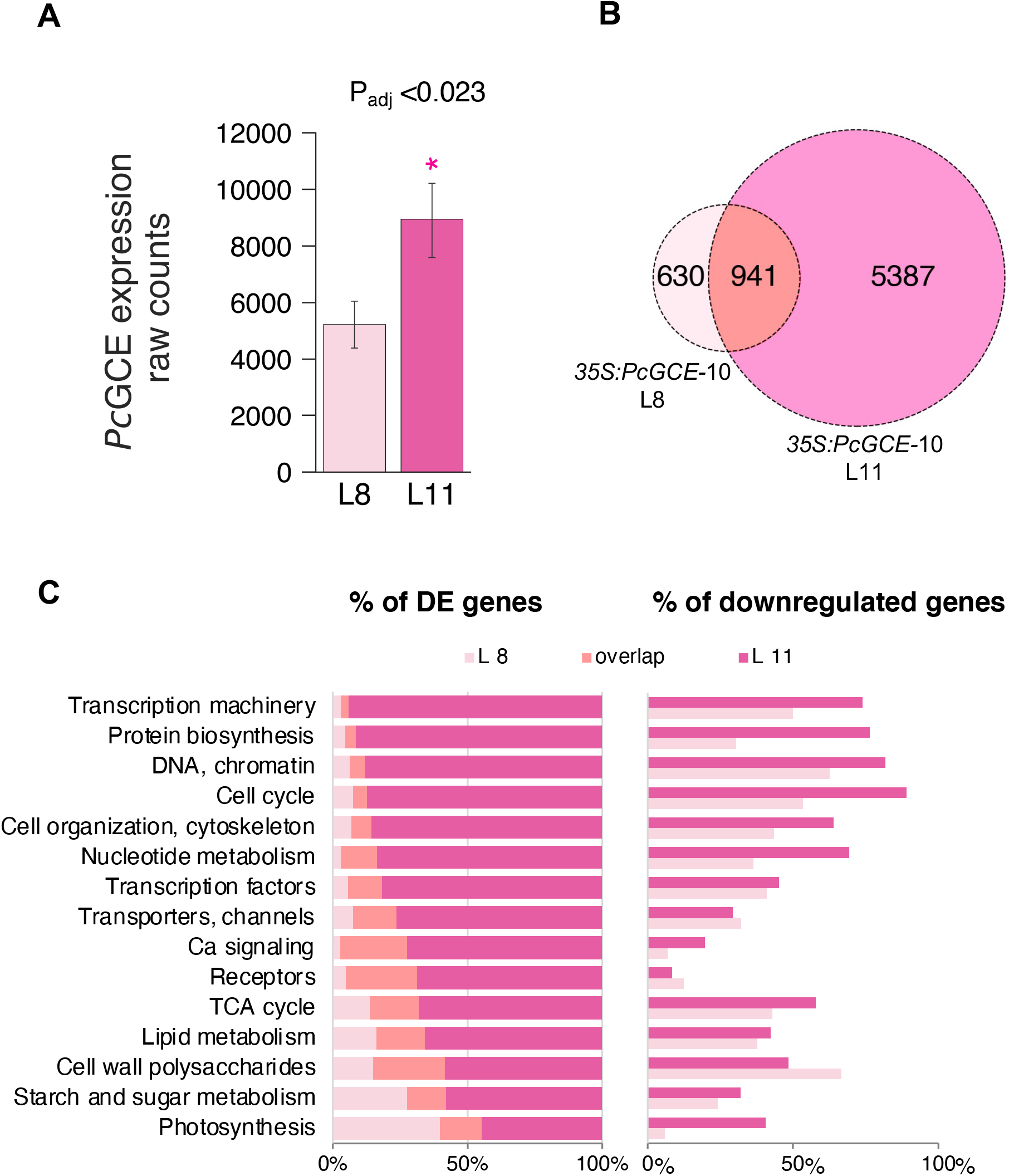
Overview of transcriptomic changes in young developing aspen leaves of *35S:PcGCE line 10*. (A) Differential expression analysis of the transgene between leaf 11 (L11) and 8 (L8) reveled significantly higher levels of *PcGCE* transcripts in L11 (Log_2_FC=0.68, P_adj_ <0.023). (B) Venn diagram of differentially expressed genes [DEGs: P_adj_<0.05 and abs(Log_2_FC) > 0.3] in transgenic compared to WT plants. C) Proportions of DE genes in L8, both L8 and L11, and L11 (left) and proportion of downregulated genes in each leaf (right) by different functional groups. Shown are examples of functional groups of DE genes. All DE genes are listed in Table S7. First unfolded leaf (L8) and young expanding leaf (L11) were analyzed in five trees.

To further reveal *Pc*GCE perception-related genes, we focused on the early upregulated genes encoding receptors and signal transduction members, including cell wall, redox, signaling, transporters and channels, stress-related hormones, and transcription factors categories (Table S9). This resulted in a list of 464 genes. To further narrow down this candidate list, we examined their expression in developing wood using Aspwood database (Sundell et al., 2017) available at Popgenie website (www.popgenie.com). We reasoned that sought candidates should not be expressed in xylem cells forming secondary cell walls, as indicated by lack of biotic stress responses in *WP:PcGCE* lines (Fig. 6). This filtering resulted in identification of 118 first sort candidates which were not expressed in wood developing tissues, and additional 276 of second sort candidates expressed in other stem tissues than secondary-wall forming xylem (Table S9). Majority of selected genes were upregulated in leaf 11 as well, and their degree of upregulation was by large higher for the candidates of sort one. We further analyzed the expression of the sort one and sort two candidates in different aspen tissues using public databases provided by Sundell et al., (2015) and Immanen et al. (2016). Majority of these genes exhibited highest expression in leaves, especially the leaves subjected to biotic and abiotic stress (Fig. 8). This indicates that the identified candidate genes are likely to mediate stress responses in leaves.

**Figure 8.**
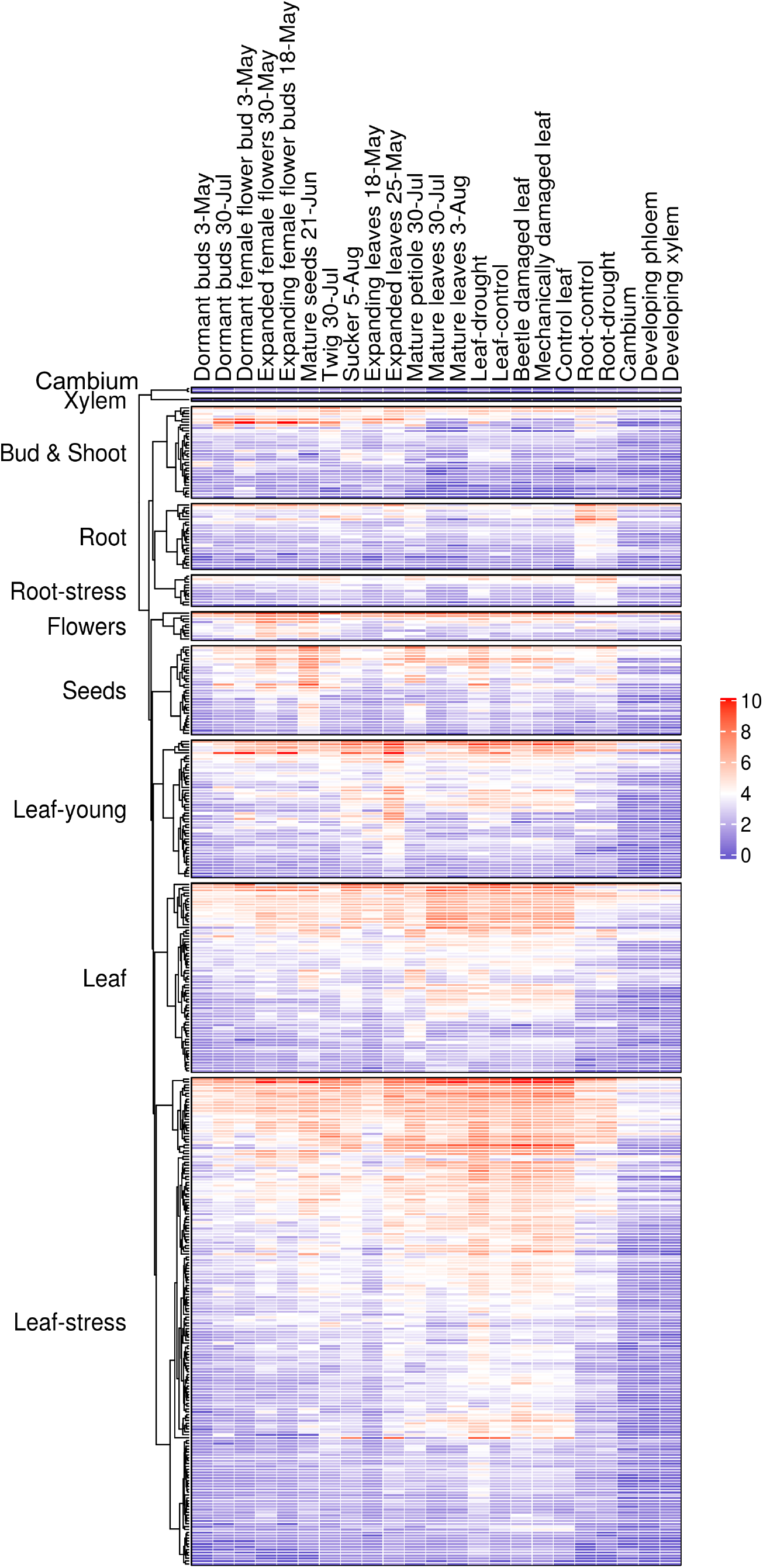
Expression patterns of selected candidates for early *Pc*GCE perception and signaling. The candidates (Table S8) were selected based on assigned putative function, upregulation in leaf 8 relative to WT and no expression in secondary wall forming xylem (Sundell et al., 2017). Normalized expression in different tissues based on expression databases described by Sundell et al. (2015) and Immanen et al. (2016) processed as described by Kumar et al.,(2020).

The identified candidate lists included, among others, several genes encoding cell surface immune receptors, receptor-like proteins (RLPs), receptor-like kinases (RLKs), wall associated kinases (WAKs), malectin domain (MD) proteins, and glutamate receptors (GLR) (Table S9). There were factors associated with ET and ABA signaling. Among transporters there were transmembrane calcium transporters, cyclic nucleotide binding ion channels (CNGCs), calcium-transporting AUTOINHIBITED Ca^2+^ ATPases (ACA), ABC channels, glutamate receptors, one MLO protein and several highly upregulated metal channels. Several pectin methylesterases (PMEs), chitinases, and callose hydrolases (GH17) were found. We also identified many transcription factors, in particular, several WRKY factors. Some of these genes were found similarly affected in Arabidopsis stems ectopically overexpressing *Pc*GCE (Tsai et al., 2017), as shown in Table S9. The identified genes constitute transcriptional modules that likely operate in recognition of *Pc*GCE and other molecular patterns associated with pathogens to activate defense systems in aspen.

### Mutated PcGCE that does not have esterase activity induces similar stress responses in aspen as the active enzyme

*Pc*GCE could be perceived in hybrid aspen leaves as a pathogen-associated molecular pattern (PAMP) or its enzymatic activity could generate oligomers signaling cell wall damage. To distinguish between these two possibilities, we generated point mutation S217A in the active site of *Pc*GCE and expressed mutated *Pc*GCE^S217A^, which lacks esterase activity in hybrid aspen under control of *35S* promoter. All four investigated transgenic lines expressing the mutated enzyme displayed leaf necrosis, premature senescence and reduced growth after 7-week-cultivation in the greenhouse (Fig. 9ABC). The severity of these symptoms correlated with the transgene expression levels (Fig. 9D), where lines with highest expression of *Pc*GCE^S217A^ exhibited most pronounced phenotypes. Glucuronoyl esterase activity was detected in proteins extracted from leaves of plants expressing wild-type *Pc*GCE whereas no activity was detected in the plants expressing mutated *Pc*GCE^S217A^ (Fig. 9F) indicating that the perception of *Pc*GCE protein is capable of triggering defense responses.

**Figure 9.**
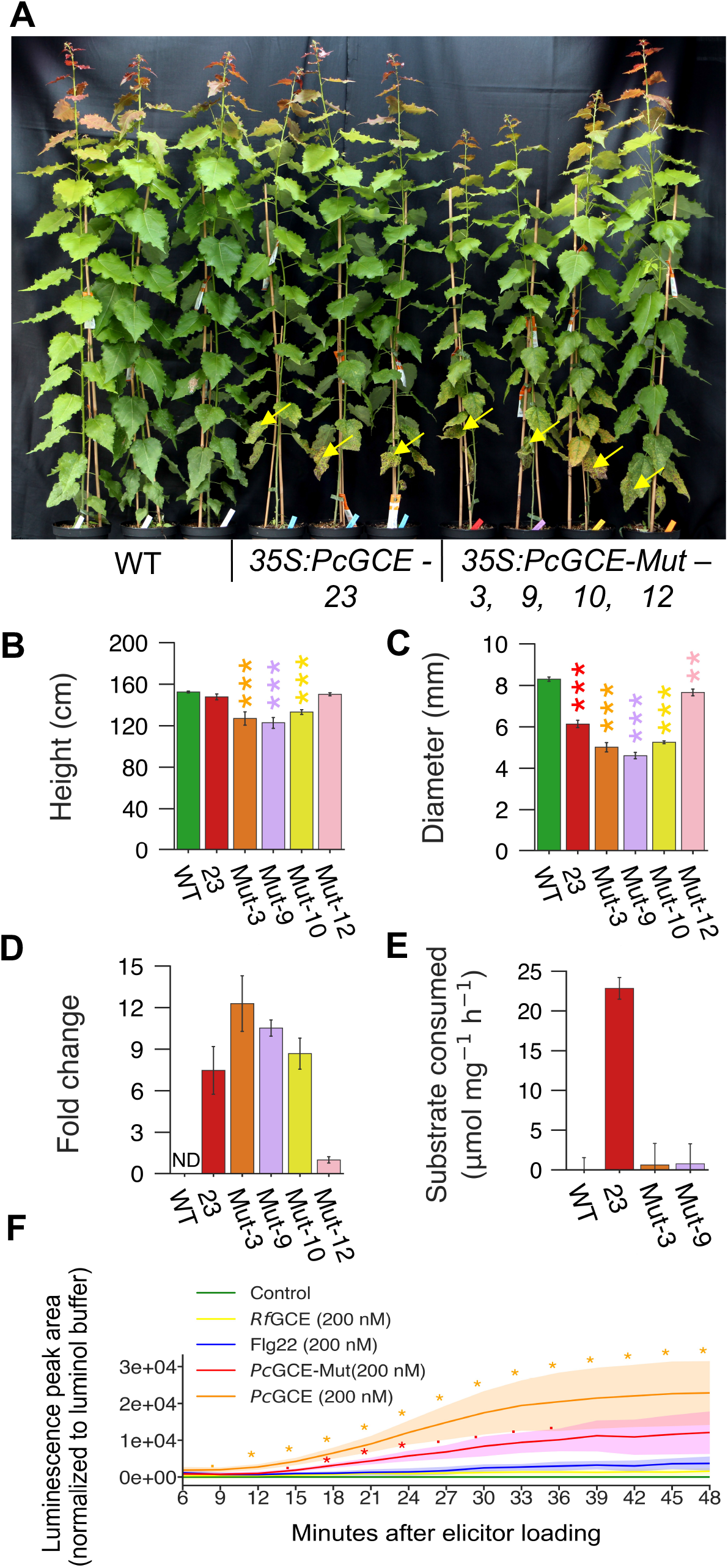
Enzymatic activity of *Pc*GCE is not needed to induce disease symptoms and stress responses in aspen. Appearance (A), height (B), diameter (C), transgene transcript levels (D), and glucuronoyl esterase activity in mature leaves necrosis-free leaves (E) of trees expressing mutated, enzymatically inactive *Pc*GCE^S217A^ from *35S* promoter (*35S:PcGCE-Mut*) as compared to WT trees and line *35S:PcGCE-23* expressing native *PcGCE*, after 7-week-cultivation in the greenhouse. Note premature leaf senescence and necrotic leaves in *35S:PcGE* and *35S:PcGCE-Mut* lines (arrows in A). (B-E) Means ± SE, n=7 (B, C) or 4 (D, E); asterisks show means significantly different from WT (Dunnett’s test; · - P≤ 0.1; * - P ≤ 0.05; ** - P ≤ 0.01; *** - P ≤ 0.001); ND-not detected. (F) Luminol-based assay for detection of elicitor-induced production of ROS in aspen leaves. Elicitor activity of *Pc*GCE and *Pc*GCE-mut compared to activity of flg22 peptide and bacterial *Ruminococcus flavefaciens* glucuronoyl esterase (*Rf*GCE). Means ± CI68%; n = 4 independent experiments. Asterisks show significant differences compared to buffer control (Dunnett’s test; · - P≤ 0.1; * - P ≤ 0.05).

Next, we expressed and purified active and mutated S217A *Pc*GCE proteins in yeast and tested their elicitor activity in aspen leaves using the luminol assay (Bisceglia et al., 2015). Aspen leaf discs were treated with either *Pc*GCE, *Pc*GCE^S217A^, bacterial glucuronoyl esterase from *Ruminococcus flavefaciens* (*Rf*GCE), flagellin-derived flg22 peptide or with buffer without elicitors. Both active and inactive *Pc*GCE induced ROS in aspen leaves, with notably higher levels than flg22 (Fig. 9F). *Rf*GCE applied to aspen leaves did not induce ROS. These results indicate that *Pc*GCE protein is sufficient for ROS induction, independently of glucuronoyl esterase activity.

## Discussion

### PcGCE is a potent inducer of defense responses in plants

*P. carnosa* is a basidiomycete pathogen of forest trees found on bark and wood of conifers (Burt, 1925). It decomposes wood causing white rot is considered as a source of potent lignocellulolytic enzymes with a wide range of technological applications (MacDonald et al., 2011; Suzuki et al., 2012). Among other enzymes, a CE15 family enzyme glucuronoyl esterase is predicted to reduce lignocellulose recalcitrance to saccharification caused by lignin carbohydrate cross-links (Bååth et al., 2016). Indeed, ectopically expressing *Pc*GCE in plants improved xylan extractability in *A. thaliana* (Tsai et al., 2012) and cellulose conversion to glucose in aspen (Gandla et al., 2015). However, it also induced premature leaf senescence and impaired growth. Here we present a detailed analysis of aspen off-target responses to *Pc*GCE propose their mechanisms and demonstrate an alternative strategy for deployment of this enzyme that avoids such off-target effects.

Reaction to ectopically expressed *Pc*GCE in aspen included inhibition of stem diameter and height growth, and very strongly suppression of root system development. Detailed analysis of leaves of transgenic plants revealed altered leaf expansion, defective hydraulic conductivity of xylem due to blockage by tyloses and gels, increased formation of calcium oxalate crystals, presence of necrosis and premature shedding (Fig. 1). These alterations were gradually developed as leaves matured. Transcriptomes of developing leaves were massively altered even at youngest developmental stages long before any visible disease symptoms appeared (Fig. 4, 7; Table S7). These early changes in transcriptome indicated stimulated development of photosynthetic machinery, altered sugar and lipid metabolism and activation of signaling of biotic stress pathways involving JA and ET. At later developmental stages, we additionally detected an activation of SAR as evidenced by increase in ROS content and *RBOHD* expression (Fig. 2, 4), increase in SA content (Fig. 3) and activation of SA-related transcriptome (Fig. 4; Table S3, S7). These responses are typical to pattern-triggered immunity (PTI) induced be various pathogen- or damaged plant-derived molecules during pathogen attack (Yu et al., 2017).

Similar to aspen, Arabidopsis expressing *Pc*GCE exhibited transcriptomic changes typical of biotic stress responses, SA signaling (Tsai et al., 2017). Moreover, several biotic-stress related genes were affected in common in both species, including glutamate receptor GLR2.7, recently shown to constitute a core stress response gene (Bjornson et al., 2021), WRKY51 that mediates JA signaling inhibition by SA and low oleic acid levels (Gao et al., 2011), SOBIR1 that together with BAK1 and RLK23 recognizes NLP20 peptide inducing necrosis (Albert et al., 2015), and other genes which we identified as putative candidates for early *Pc*GCE responses in aspen leaves (Table S9). This suggests that a common perception system for *Pc*GCE protein exists in diverse plants.

### PcGCE mRNA is not cell-to-cell mobile, increases with leaf age and induces systemic stress signals

Grafting experiments showed that *Pc*GCE mRNA is only present in transgenic plant parts (Fig. 5) indicating that the transcripts are not cell-to-cell mobile. Unexpectedly, the level of transgene transcripts increased exponentially with leaf age (Fig. 4), which must be due either to age-dependent increase in *35S* promoter activity or to high stability of *Pc*GCE transcripts resulting in their accumulation. The cauliflower mosaic virus (CaMV) *35S* promoter includes several response elements such as As-1, As-2, Dof and I-box that provide means for regulated activity in plants (Bhullar et al., 2007) and there are reports of *35S*-driven transgene induction in response to heat (Boyko et al., 2010) or aging (Kiselev et al., 2021). There were several transcripts corresponding to heat shock- and senescence-related proteins in *35S:PcGCE* lines (Table S3, S7), suggesting that *Pc*GCE induces activity of 35S promoter, thus creating a positive regulatory feedback loop resulting in exponential transgene accumulation.

Grafting experiments showed that some effects of *Pc*GCE on gene expression are systemically induced and depend on the presence of distant transgenic leaves where the transgene is expressed (Fig. 5). Such leaves are likely a source of diffusible signal(s) inducing systemic responses. In contrast, systemic increase in ROS content in WT shoots was observed also in the absence of transgenic distal leaves (Fig. 5). This rises a possibility that systemic ROS signals could be generated in a different way than systemic signals involved in gene regulation. Induction of ROS but not marker gene expression was observed from developing xylem cells expressing *Pc*GCE (Fig. 6), suggesting that developing xylem cells of transgenic rootstocks could be a source of ROS signals in WT scions. Propagation of such signals might involve H_2_O_2_ wave along veins (Choi et al., 2016; Lew et al., 2020).

### PcGCE protein is perceived as PAMP and involves upregulation of large number of proteins

PTI -like responses to ectopically expressed *Pc*GCE were observed in transgenic aspen plants expressing mutated form of *Pc*GCE that is enzymatically inactive (Fig. 9), indicating that *Pc*GCE protein is recognized as PAMP. The exact sequence that is recognized by plant is currently unknown. We did not find significant similarity between *Pc*GCE AA sequence and known elicitor peptides such as peptides from EIXs (Noda et al., 2010; Frías et al., 2019), flg22 (Naito et al., 2008), elf18 (Kunze et al., 2004) or nlp20 (Böhm et al., 2014b). Nevertheless, *Pc*GCE is a new microbial enzyme with demonstrated elicitation of defense responses in so diverse plants as aspen and Arabidopsis, suggesting a broad specificity of epitope recognition. Since its elicitor activity is independent of its enzymatic activity, it is possible that other glucuronoyl esterases from other microorganisms do not show such elicitation. This is supported by a lack of ROS induction by exogenously applied bacterial *Rf*GCE to aspen leaves (Fig. 9F).

Receptor(s) for *Pc*GCE epitope are currently unknown, but the transcriptomics analysis of early developmental leaf stages provides large number of potential candidates. A striking results of transcriptome analyses (Table S3, S7, S9) is an evidence for regulation of i) multitude of signal transduction modules and ii) regulation of groups of similar potential members of these modules by *Pc*GCE. This on one hand reflects complexity of immune responses as recently discussed by Kanyuka and Rudd (2019) and on the other is in line with current concepts of a need for redundancy in the pathogen perception and response systems to allow rapid evolution (Adachi et al., 2019). Among the affected genes, we find several members of the modules for pattern recognition described in other plant species (Böhm et al., 2014a; Couto and Zipfel 2016; Yu et al., 2017; He et al., 2018; Kanyuka and Rudd 2019). Because it is secreted to the apoplast, *Pc*GCE protein is expectedly recognized by a surface PRR. Coordinated up-regulation of cell surface LRR-RLPs and LRR-RLKs suggests the involvement of these proteins and their complexes. An example of a complex that could be involved is SOBIR1-RLP-BAK1 complex (Jehle et al., 2013; Zhang et al., 2013; Albert et al., 2015; Zhang et al., 2014). RLP recognizing *Pc*GCE could be RLP13 (*Potri.001G064100*), RLP21 (*Potri.006G061000*), RLP27 (*Potri.T164200* and *Potri.T114000*), RLP34 (*Potri.T086100*), RLP41 (*Potri.T164100*) or RLP43 (*Potri.012G005600*). Some of these RLPs are very similar, and likely have redundant functions. BAK1 function could be encoded by *Potri.003G023000* or *Potri.001G206700* (Table S7) and SOBIR1 by *Potri.012G090500* or *Potri.015G086800* (Table S9). The SOBIR1-RLP43-BAK1 model is supported by induction of SOBIR1, RLP43, and BAK1 in *Pc*GCE-expressing Arabidopsis (Tsai et al., 2017). Members of the PRR complexes have been shown to rapidly trans-phosphorylate each other within the complex and induce transient increase in intracellular calcium via passive or ATP-driven channels (Yu et al., 2017), several of which, including glutamate receptors, were identified upregulated at early leaf developmental stages (Table S7, S9). Further, the extracellular alkalization and ROS burst is typically observed (Yu et al., 2017). A key player in the ROS is RBOHD (*Potri.003G159800*) along with extracellular peroxidases, and these enzymes were highly upregulated in leaf 9 and 11 (Table S7, S9).

The transcriptomics revealed many other potential receptors, which are likely involved in downstream pathways triggered by the *Pc*GCE perception. One example is *FLS2* (*Potri.004G065400*). In Arabidopsis, FLS2 together with BAK1 perceives flagellin-derived peptide fl22 providing resistance to a broad range of bacterial pathogens (Chinchilla et al., 2007). *FLS2* was also induced in *Pc*GCE-expressing Arabidopsis (Tsai et al., 2017). Another example is RALF-mediated signaling. Two *RALF27* homologs (*Potri.017G098100* and *Potri.017G098300*) were upregulated by *Pc*GCE (Table S9). *RALF27* homologs were found in genomes of poplar fungal pathogens *Sphaerulina musiva* (Peck) Quaedvlieg, Verkley & Crous and *Septoria populicola* Peck but their role in virulence of these pathogens is unclear (Thynne et al., 2017). Nevertheless, the finding suggests a role for RALF27 in aspen immune responses. Incidentally, we observed upregulation of two clade XII malectin domain (MD) aspen genes, *PtMD121* and *PtMD105* (Kumar et al., 2020), which could be involved in RALF27-mediated signaling, as reported for other members of *Cr*RLK1L family (Zhang et al., 2020). *Pc*GCE induced genes also included *ZAR1* homologs (*Potri.006G014400*, *Potri.005G119800*, and *Potri.016G092600*) and *ZAR1* was similarly upregulated in Arabidopsis (Tsai et al., 2017). ZAR1 is an NLR type of resistance protein forming the resistome complex in response to several effectors and mediating rapid HR (Adachi et al., 2019). We also observed several genes associated with pectin modification, including *PtPME2*, *PtPME3* and other pectin methyl esterases, upregulated early during leaf development in transgenic plants (Table S9). At a later developmental stage (leaf 11), genes encoding polygalacturonases (PGs) were upregulated whereas several pectate lyases from family PL1 were downregulated (Table S7). PGs could be responsible for release of oligogalacturonides (OGs) to the apoplastic space, which are perceived by wall associated kinases (WAKs) activating a wide range of defense responses (Ferrari et al., 2013, Kohorn, 2016). *WAK2* homologs (*Potri.002G076000* and *Potri.002G075900*), and homologs of *WAKL1* (*Potri.009G154600*) and *WAKL5* (*Potri.009G154600*) were highly upregulated in aspen by *Pc*GCE (Table S9) and *WAK2* was also upregulated by this enzyme in Arabidopsis (Tsai et al., 2017). WAK2 mediates immune responses via mitogen-activated protein kinase 6 (MPK6) (Kohorn et al., 2009). These examples show that *Pc*GCE induces a series of immune responses possibly inducing defenses to pathogens.

### Specificity of transgene expression provides a powerful mean to avoid detrimental responses to transgene-encoded proteins

We showed that expressing *Pc*GCE from WP promoter derived from aspen *GT43B* gene, which is active in cells depositing xylan-type secondary walls (Ratke et al., 2015) avoids growth inhibition and premature senescence phenotypes that are observed in plants ectopically expressing the transgene (Fig. 6). The following observations indicate that this is due to a lack of *Pc*GCE perception in secondary wall forming cells: i) majority of genes that are upregulated by *Pc*GCE highly expressed in leaves and are not or only weakly expressed in the developing wood tissues (Fig. 8); ii) transcripts of *Potri.012G005600* encoding RLP43 which is the prime candidate for *Pc*GCE perception are not detected in developing wood (Table S9); iii) systemic induction of immune marker gene expression required expression of *Pc*GCE in the leaves whereas its expression in the stems or roots was not capable of inducing systemic signals (Fig. 5). These observations suggest that systemic immune responses of *Pc*GCE are initiated in the leaves. Since main pathogen entry is via epidermis or stomata and this is where the perception machinery is expectedly expressed, directing the expression of transgenes of microbial origin away from leaf epidermis and mesophyll can be a good strategy to avoid immune responses in general.

## Experimental Procedures

### Generation of transgenic aspen lines

*Phanareochete carnosa* Burt glucuronoyl esterase (*Pc*GCE) cDNA (NCBI accession: JQ972915; Tsai et al., 2012) with its native signal peptide replaced by the signal peptide of *Ptt*Cel9B3 (GenBank accession AY660968.1) as described previously (Gandla et al, 2015) was used for the cloning of *WP:PcGCE* construct. The transgene in pENTR/D-TOPO vector (pENTR™ Directional TOPO Cloning Kits, Invitrogen) was subcloned into vector pK-pGT43B-GW7 (Ratke et al., 2015) containing the *WP* promoter.

To clone the vector *35S:PcGCE*^S217A^ with *Pc*GCE carrying a point mutation S217A in the active site, pENTR/D-TOPO vector (pENTR™ Directional TOPO® Cloning Kits, Invitrogen), containing *Pc*GCE cDNA with exchanged native signal peptide was PCR amplified to introduce a point mutation *Pc*GCE^S217A^ using overlapping primers (Table S10). The pENTR/D-TOPO vector product was subsequently subcloned into binary vector pK2WG7.0 (Karimi et al., 2002) using Gateway® System (Invitrogen).

Vectors were transferred into competent *Agrobacterium tumefaciens* (Smith and Townsend) Conn strain GV3101 by electroporation and used to transform hybrid aspen, *Populus tremula* L. x *tremuloides* Michx., clone T89 as described in Gandla et al. (2015). Twenty independent lines were obtained and clonally propagated and three lines of each construct with the highest expression levels were selected for this study.

### Glucuronoyl esterase activity

Total proteins were extracted as previously described by Gandla et al. (2015) from mature necrosis-free leaves. Shortly, 100 mg of frozen powder material was added to 0.5 mL of extraction buffer [50 mM sodium phosphate buffer pH 6.0, 2 mM EDTA, 4% PVP mw 360,000, 1 M NaCl, and protease inhibitor cocktail (cOmplete, Roche)], and the sample was incubated for 1 h at 4°C. After centrifugation (20,000g, 15 min), the supernatant was desalted and concentrated using a column (Nanosep 30k omega OD30C34) by centrifugation (4,000g, approx. 10 min, four times) with four washes of 50 mM sodium phosphate buffer. Glucuronoyl esterase activity was measured using benzyl-D-glucuronate (Santa Cruz Biotechnology, sc-221344) as a substrate. The reaction mix having a final volume of 40 μL containing 50 mM sodium phosphate buffer pH 6, 5 mM substrate and extract containing 5μg of protein extract, was incubated at 30°C for 60 min. Remaining substrate was quantified by adding 80 μL of alkaline hydroxylamine hydrochloride (2 M hydroxylamine hydrochloride and 3.5 N sodium hydroxide, 1:1, v:v), incubating for 5 min, acidifying with 40 μL of 12% hydrochloric acid and adding 40 μL of 0.37 M ferric chloride before absorbance measurement at 540 nm (Hestrin, 1949). The standard curve was generated using between 0.1 mM to 3 mM benzyl-D-glucuronate.

### Analysis of calcium oxalate crystals

Successive leaves were collected from transgenic *35S:PcGCE* plants (lines 4 and 10) and WT. They were stored in 70% ethanol before clearing in 2.5% commercial bleach solution of sodium hypochloride (NaClO) for approximately 8 hours until the chlorophyll was removed. Small sections from top, middle and bottom of each leaf were processed in four trees per genotype. The sections were washed with water and mounted in 50% glycerol for the observation using Axioplan 2 microscope (Zeiss, Germany) and Nomarsky’s optic. Counts per vein area were performed using Image J, and data for top, middle, and bottom of the leaf were averaged.

Crystal chemical nature was determined by X-ray diffraction (XRD) using ethanol-fixed dissected veins from leaf 20 of transgenic (35S:GE line 10) and WT plants. The XRD analysis was performed by a Bruker D8 advance X-ray diffractometer with Cu Kα radiation in ϴ:ϴ mode, equipped with a super speed VÅNTEC-1 detector. The samples were directly mounted on a Si low-background rotating sample holder and analyzed by continuous scanning for at least 4 h. Crystals were identified using Bruker software and the powder diffraction file PDF-2 (International Center for Diffraction Data, Newtown Square, USA, 2010).

### Dye uptake experiments

To monitor hydraulic continuity of xylem in leaves, transgenic (two independent lines) and WT small lateral branches were placed in tubes containing 4 % basic fuchsin for 3-4 h after which the leaves were photographed with illumination from above.

### DAB staining

1 cm^2^ leaf pieces from freshly collected leaves were submerged in 1 ml Diaminobenzidine tetrahydrochloride (DAB) reagent solution (1 mg/ml), pH 7.0 and incubated for ~ 24h at room temperature in the dark. Then the DAB solution was replaced with water followed by clearing the leaves in 95% ethanol for 24 h at 37 °C, rehydrating using a graded ethanol series and storing in 0.05 M phosphate buffer pH 6.5. The material was mounted in 50% aqueous glycerin and observed using Axioplan 2 Universal Microscope (Carl Zeiss, Jena, Germany). The micrographs were converted to grey scale with pixel intensity between 0 and 256 using Pythons library OpenCV (https://pypi.python.org/pypi/opencv-python). The area of dark pixels (grey scale 0-31) considered as DAB signals was calculated as % of total area.

### Microscopy of cuticle

Cuticle was analyzed in fully expanded leaves in three trees of each of two independent transgenic lines per construct and WT. The leaf pieces were fixed in 2.5% glutaraldehyde, embedded in Steedman’s wax, and stained for lipids with Nile red (Sigma-Aldrich) as described by Dobrowolska et al. (2015). Sections were examined by epifluorescence microscopy (Axioplan 2; Zeiss, Germany) with excitation filter 450–490 nm and barrier filter 520 nm. Cuticle thickness was measured using Image J in six sections per tree, at six random places per section, for each abaxial and adaxial cuticle.

### Leaf wax and fatty acid analysis by fatty acid methyl esters (FAMEs)method

For wax analysis, a 2 cm-diameter leaf disc was immersed in 1 mL chloroform and shaken for 30 sec, and the total solvent volume was transferred into a micro vial. The chloroform volume was evaporated by flushing with N_2_ gas, until a wax pellet remained, and the material was stored at −80°C until analysis. The waxes were derivatized by methoxyamination and silylation as described in Gullberg et al. (2004). The derivatized samples were analyzed as described in Diab et al. (2019b). The wax content is expressed as relative concentration.

For fatty acid analysis, 19 to 21.6 mg of ground frozen leaf material was extracted with 500 μl of extraction buffer [2:1 v/v chloroform (Darmstadt, Germany): methanol (Waltham, MA, USA)] including following a modified Folch’s protocol (Diab et al., 2019a). The fatty acids were converted into their corresponding Fatty Acid Methyl Esters (FAME) by methylation with diazomethane. For absolute quantification, a calibration curve was prepared from the commercially available Supelco 37 Component FAME Mix at 7 levels. Analysis was performed by GC-QqQ-MS equipped with a Zebron ZB-FAME 20 m x 0.18 mm i.d. fused silica capillary column.

### Data analysis /statistical methods for metabolomics

For the GC-MS data, all non-processed MS-files from the metabolic analysis were exported from the ChromaTOF software in NetCDF format to MATLAB R2016a (Mathworks, Natick, MA, USA), where all data pre-treatment procedures, such as base-line correction, chromatogram alignment, data compression and Multivariate Curve Resolution were performed using custom scripts. The extracted mass spectra were identified by comparisons of their retention index and mass spectra with libraries of retention time indices and mass spectra 3. Mass spectra and retention index comparison was performed using NIST MS 2.0 software. Annotation of mass spectra was based on reverse and forward searches in the library. Masses and ratio between masses indicative of a derivatized metabolite were especially notified. If the mass spectrum according to SMC’s experience was with the highest probability indicative of a metabolite and the retention index between the sample and library for the suggested metabolite was ± 5 (usually less than 3) the deconvoluted “peak” was annotated as an identification of a metabolite.

Principal component analysis (PCA) was performed in the R (v3.4.0; R Core Team, 2019) program, by using ggfortify (Horikoshi and Tang, 2018) package.

### Transcriptomic analyses

Developing leaves (leaf number 8, 11, 21 and 23) were collected from ten-week-old hybrid aspen grown in the greenhouse. The RNA was extracted as described in Ratke et al. (2018). Five biological replicates of leaves 8, and 11 of line *35S:PcGCE - 10* and corresponding WT; as well as four biological replicates of leaves 21 and 23 of lines *35S:PcGCE* - 10, *35S:PcGCE* - 23, *WP:PcGCE* - 8, *WP:PcGCE* - 14 and eight biological replicates of corresponding WT were analyzed. cDNA was sequenced using Illumina HiSeq-PE150 platforms of Novogene Bioinformatics Technology Co., Ltd. (Beijing). Quality control and mapping to *P. trichocarpa* transcriptome v3.0 of leaf 8 and 11 samples were performed by Novogene. For other samples, RNA-seq raw data were filtered and mapped using methods described by Kumar et al. (2020). Raw counts were used for differential expression analyses using DESeq2, comparing genotype *35S:PcGCE*-10 (leaf 8, 11, 21 and 23), *35S:PcGCE*-23 (leaf 21and 23), *WP:PcGCE*-8 (leaf 21 and 23) and *WP:PcGCE*-14 to WT. The P-values were corrected for multiple testing using the Benjamini and Hochberg method to calculate P_adj_. Genes were considered as DE when P_adj_ < 0.05 and abs(Log_2_FC) > 0.3. For further processing of the data such as gene sorting, filtering, intersection, sample grouping and biological function summary, R (v3.4.0; R Core Team, 2019) and Python (Van Rossum and Drake, 2009) programs were used.

### Reverse transcription-quantitative polymerase chain reaction

Developing leaves and developing xylem tissues scrapped from the stems after bark peeling were used for RNA extractions as described in Ratke et al. (2015) using three or four biological replicates. cDNA was synthesized using Bio-Rad iScript™ cDNA Synthesis kit. Quantitative polymerase chain reactions (qPCRs) were performed using LIGHTCYCLER 480 SYBR GREEN I Master Mix (Roche) either using 20 μL reaction volume in a LightCycler® 480 System II (Roche), or using 5 μL reaction volume in a C1000 Touch thermal cycler (Bio-Rad). The PCR program was: 95°C for 5 min, then 50 cycles of 95°C for 30 s, 60°C for 15 s, and 72°C for 30 s. *UBQ-L* (*Potri.005G198700*) was selected as reference gene based on GeNorm (Vandesompele et al., 2002) reference gene analysis chosen from four different genes. Primers are listed in Table S10. The relative expression level was calculated according to Pfaffl (2001), in the Python programming language (Van Rossum and Drake, 2009).

### Grafting

The grafts were prepared from 5-week-old WT, *35S:PcGCE*-10 and *35S:PcGCE*-23 plants. The scions excised 5-10 cm below the shoot apex had their stems trimmed into wedges, inserted into longitudinally cut rootstocks between the stem halves, and sealed with Parafilm (Pechiney Plastic Packaging Company). Upper part of shoots of the grafted plants were enclosed in transparent plastic bags covering them from 10 cm bellow the stock/scion interface to the apex of the plant. Leaves were removed from the rootstocks of half of the grafted plants at the day of grafting. The bags were removed 7 to 10 days after grafting.

### Hormonomics

Leaves 21 and 23 of four biological replicates of lines *35S:GCE-10* and *35S:GCE*-23 and eight biological replicates of WT were ground in liquid N_2_. Samples were extracted, purified and analyzed according to method described in Šimura et al., (2018). Mass spectrometry analysis of targeted compounds (Table S11) was performed by an UHPLC-ESI-MS/MS system comprising of a 1290 Infinity Binary LC System coupled to a 6490 Triple Quad LC/MS System with Jet Stream and Dual Ion Funnel technologies (Agilent Technologies, Santa Clara, CA, USA). The quantification was carried out in Agilent MassHunter Workstation Software Quantitative (Agilent Technologies, Santa Clara, CA, USA).

### Metabolic profiling

Leaves 21 and 23 of transgenic and WT plants were ground in liquid N_2_ and 9 - 12 mg portions were extracted in 500 μL of extraction buffer (20/20/60 v/v chloroform (Darmstadt, Germany): water Milli-Q: methanol (Waltham, MA, USA)) including internal standards (Gullberg et al., 2004). The standards: L-proline-^13^C5, alpha-ketoglutarate-^13^C4, myristic acid-^13^C3, and cholesterol-D7 (Andover, MA, USA), as well as succinic acid-D4, salicylic acid-D6, L-glutamic acid-^13^C5,15N, putrescine-D4, hexadecanoic acid-^13^C4, D-glucose-^13^C6, D-sucrose-^13^C12 (Sigma, St. Louis, MO, USA), were added to ground frozen leaf material. The sample was shaken with a tungsten bead in a mixer mill at 30 Hz for 3 minutes, the bead was removed and the sample was centrifuged at +4 °C, 20 000 g, for 10 minutes. 75μL of supernatant was transferred to a micro vial and solvents were evaporated. Small aliquots of the remaining supernatants were pooled and used to create quality control (QC) samples. The samples were analyzed in batches according to a randomized run order on GC-MS.

Derivatization and GC-MS analysis were performed as described previously (Gullberg et al. 2004). 0.5 μL of the derivatized sample was injected in splitless mode by a L-PAL3 autosampler (CTC Analytics AG, Switzerland) into an Agilent 7890B gas chromatograph equipped with a 10 m x 0.18 mm fused silica capillary column with a chemically bonded 0.18 μm Rxi-5 Sil MS stationary phase (Restek Corporation, U.S.) The injector temperature was 270 °C, the purge flow rate was 20 mL min^-1^ and the purge was turned on after 60 s. The gas flow rate through the column was 1 mL min^-1^, the column temperature was held at 70°C for 2 min, then increased at a rate of 40°C min^-1^ to 320°C, and maintained for 2 min. The column effluent was introduced into the ion source of a Pegasus BT time-of-flight mass spectrometer, GC/TOFMS (Leco Corp., St Joseph, MI, USA). The transfer line and the ion source temperatures were 250°C and 200°C, respectively. Ions were generated by a 70 eV electron beam at an ionization current of 2.0 mA, and 30 spectra s^-1^ were recorded in the mass range m/z 50 - 800. The acceleration voltage was turned on after a solvent delay of 150 s. The detector voltage was 1800-2300 V. Data were analyzed by the same procedures as wax and fatty acids data.

### Purification of of Pc*GCE* expressed in P. pastoris

Generation of transgenic *Pitchia pastoris* (Guillermond) Pfaff strain GS115 expressing native *Pc*GCE was described previously (Tsai et al., 2012). Similar strategy was followed to express mutated *Pc*GCE^S217A^. The strains were grown for 24 h in non-inductive buffered glycerol complex medium [BMGY: 1% yeast extract, 2% peptone, 100 mM potassium phosphate (pH 6.0), 1 % yeast nitrogen base with ammonium sulfate (YNB) without amino acids, 4×10^-5^ % biotin, 1% glycerol], followed by 4 days in inductive buffered methanol complex medium (BMMY) obtained by adding 100 % methanol in BMGY to a final concentration of 0.5% methanol and then repeating the additions of the same volume every 24 h to maintain the induction during 72 h (Tsai et al., 2012). *P. pastoris* culture medium containing the recombinant protein was purified by affinity chromatography Ni-NTA agarose (Roche).

### Luminol assay for ROS

Leaf discs from young, expanded leaves of 7-week old hybrid aspen plants were sampled with a biopsy punch (4 mm diameter) and loaded into wells of 96-well plate (Roche LightCycler® 480 System) containing 100 μL of water (one sample per well). The plate was kept overnight at room temperature in the dark. Then, the water was replaced with 100 μL of assay solution containing 50mM phosphate buffer pH 6.5, 50 μM L-012 (Wako Chemicals, 120-04891), 10 μg/mL peroxidase from horseradish (Boehringer-Mannheim) and 200 nM of one of the tested elicitors: bacterial *Ruminococcus flavefaciens* glucuronoyl esterase (*Rf*GCE) (Megazyme 9016-18-6), flg22 (AnaSpec AS-62633), purified native *Pc*GCE, purified mutated *Pc*GCE^S217A^ or no elicitor (control). The assay plate light emission was measured immediately using high resolution photon counting system (HRPCS5 PHOTEK). Luminescence peak area was calculated with Image32 (6.0.3, Photek, East Sussex, UK).

## Acknowledgements

The authors are grateful to Swedish Metabolomic Centre in Umeå for carrying out metabolite analyses.

## Author Contribution

END performed majority of experiments (gene expression, transcriptomics, metabolomics, grafting, phenotyping, luminol assay) and wrote the manuscript, MDM created transgenic aspen lines and supervised phenotyping and vector cloning, XKL performed initial phenotyping, HCB performed RT-qPCR analyses, ID performed microscopy of leaves, MT and DB analyzed crystals, JL discovered tyloses and gels and performed dye uptake experiments, JS and KL performed hormonomics, LAK supervised enzymatic analyses, MEE supervised luminol assay, AT and ERM cloned and expressed *Pc*GCE in *Pichia*, and designed the active site mutation, EJM designed and coordinated the study.

## Supporting Information

**Figure S1.** Leaves of transgenic aspen expressing *35S:PcCGE* display altered free fatty acids and epicuticular wax profile.

**Figure S2.** Leaves of transgenic aspen ectopically expressing *Pc*CGE display increased deposition of prismatic crystals identified as calcium oxalate along the veins.

**Table S1.** Relative content of wax compounds extracted by chloroform form expanded leaves (leaf 14, 16, 18) of lines 10 and 23 constitutively expressing *Pc*GCE and WT. Data are integrated peaks of GC-MS chromatograms. Only identified compounds are shown.

**Table S2.** All differentially expressed genes in each transgenic line expressing *Pc*GCE, 10 and 23, and each leaf developmental stage, leaf 21 and leaf 23.

**Table S3.** Differentially expressed genes in both transgenic lines expressing *Pc*GCE, 10 and 23, and either leaf developmental stage, leaf 21 and leaf 23.

**Table S4.** All differentially expressed genes in each transgenic *WP:PcGCE* line, 8 and 14, and each leaf developmental stage, leaf 21 and leaf 23.

**Table S5.** Differentially abundant hormones, their degradation products and their precursors in leaves 21 and 23 of transgenic plants (*WP:PcGCE* lines 8 and 14, combined) compared to WT.

**Table S6.** Differentially abundant metabolites in leaves 21 and 23 of transgenic plants (*WP:PcGCE* lines 8 and 14, combined) compared to WT.

**Table S7**. All differentially expressed genes in transgenic line 10 ectopically expressing *Pc*GCE at two developmental stages, leaf 8 and leaf 11.

**Table S8.** GO enrichment analysis of DE genes in leaf 8 and 11 of *35S:PcGCE* line 10 plants compared to WT.

**Table S9.** All signal perception related genes upregulated at early leaf developmental stage in transgenic line 10 ectopically expressing *Pc*GCE and their expression in developing wood and other aspen tissues.

**Table S10.** Primers used in this study.

**Table S11.** Phytohormones and related compounds targeted by hormonomics analysis.

