## Supplementary material for "*Pc*GCE is a potent elicitor of defense responses in aspen": Suppl Figs

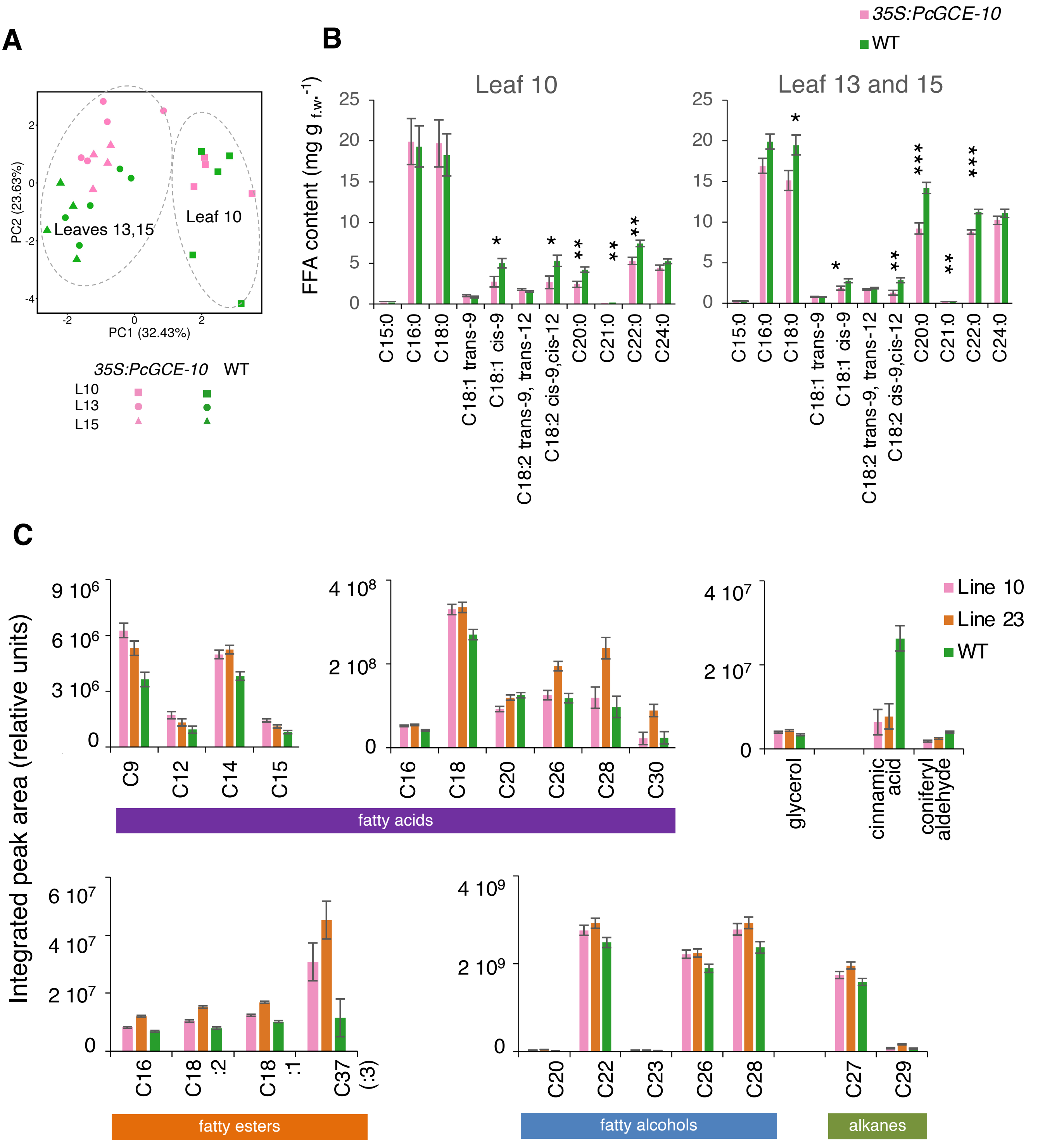


**Figure S1. Leaves of transgenic aspen ectopically expressing *PcCGE* display altered free fatty acids and epicuticular wax profile.** (A) Principal component analysis of free fatty acid (FFA) contents per fresh weight of leaves 10, 13, and 15 in transgenic (*35S:PcGCE-10*) and WT plants showing separation of expanding (leaf 10) and expanded (leaf 13 and 15) leaf samples. (B) FFA contents in expanding and expanded leaves (14, 16, 18) of transgenic and WT plants. Data are means ± SE of at least four biological replicates per line. Asterisks show significant differences between transgenic and WT plants (ANOVA Fisher's test, $P \leq 0.05$ - *; $P \leq 0.01$ - **; $P \leq 0.001$ - ***. (C) Relative contents of chloroform extracted wax compounds in expanded leaves of two transgenic lines carrying *35S:PcGCE*, line 10 and line 23, and in WT. Data are means ± SE, n=3 biological replicates, for those wax compounds that are significantly different between WT and transgenic plants (post-ANOVA contrast, $P \leq 0.05$ ). Data for other identified wax compounds are shown in Supplemental Table S1.

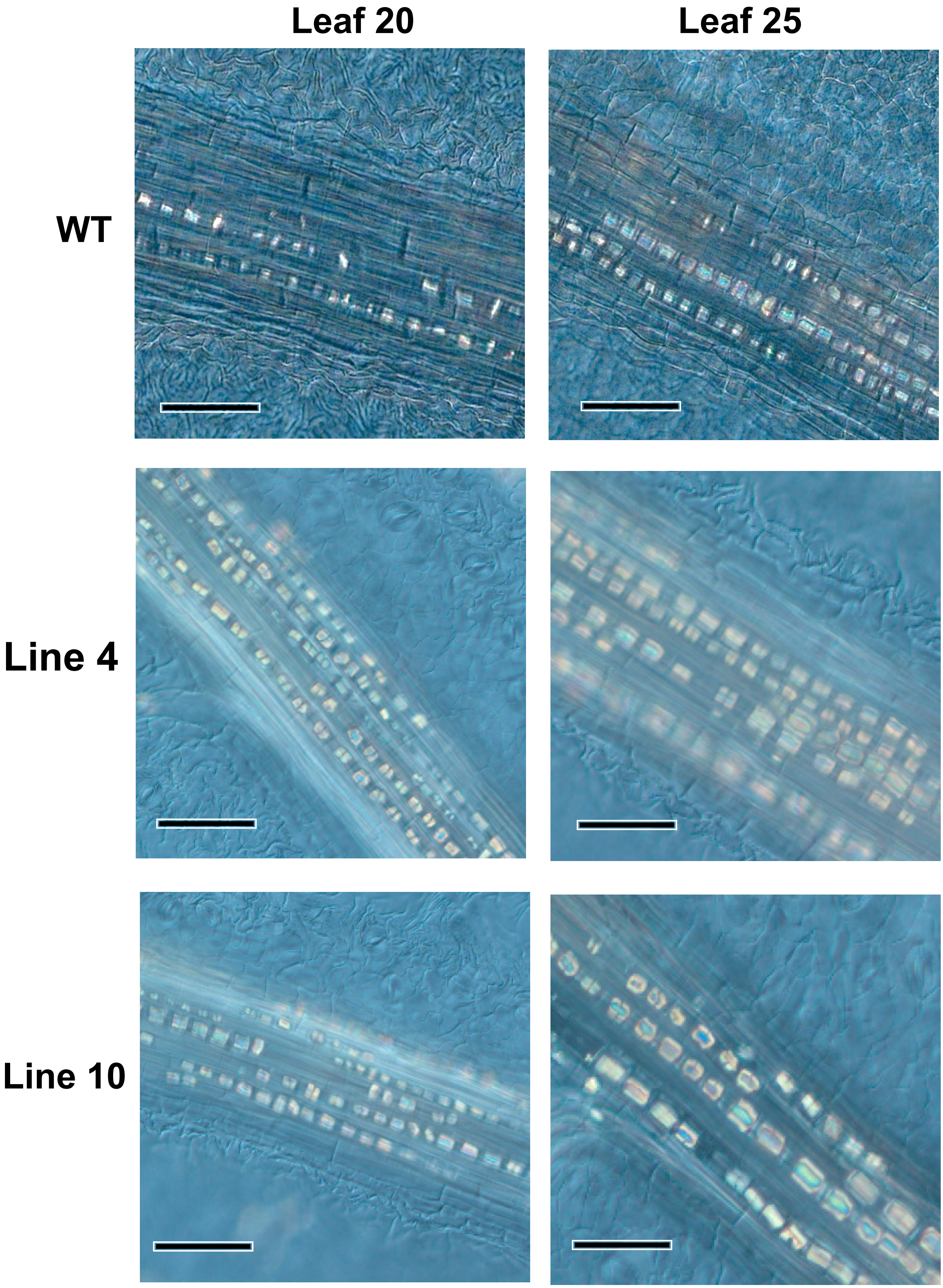


**Figure S2. Leaves of transgenic aspen expressing 35*S:PcCGE* display increased deposition of prismatic crystals along the veins identified as calcium oxalate.** Representative images of leaf 20 and leaf 25 in line 4 and 10 and WT viewed in Nomarsky's optics. Scale bars = 100 µm.
